# Neurodegeneration caused by LRRK2-G2019S requires Rab10 in select dopaminergic neurons

**DOI:** 10.1101/586073

**Authors:** Stavroula Petridi, C. Adam Middleton, Alison Fellgett, Laura Covill, Amy Stewart, Jack Munns, Friederike Elisabeth Kohrs, P. Robin Hiesinger, Laurence Wilson, Sangeeta Chawla, Christopher J. H. Elliott

## Abstract

Inherited mutations in the LRRK2 protein are the commonest known cause of Parkinson’s, but the molecular link from increased kinase activity to pathological neurodegeneration remains to be determined. *In vitro* (biochemical and cell culture) assays led to the hypothesis that several Rab GTPases might be LRRK2 substrates. Here we show that Rab10 potently modifies *LRRK2*-*G2019S* mediated electrophysiological responses in an *in vivo* screen, in which each Rab was overexpressed in *Drosophila* dopaminergic neurons. We therefore tested the effect of *Rab10* loss of function on three *LRRK2*-*G2019S* phenotypes (vision, movement and sleep) that rely on dopaminergic circuits in both flies and mammals. The knock-out of Rab10 *in vivo* fully rescues the reduced responses induced by dopaminergic *LRRK2-G2019S* in visual and motor (reaching, proboscis extension) assays, but the sleep phenotype is unaffected. We show that Rab10 is expressed in dopaminergic (tyrosine hydroxylase positive) neurons controlling vision and proboscis movement, but undetectable in those controlling sleep, indicating that anatomical and physiological patterns of Rab10 are related. Our results support the idea that LRRK2 phosphorylates separate targets in distinct neurons and confirm that one degenerative pathway starts with Rab10. Although Rab3 is another putative substrate of LRRK2, it shows no synergy with G2019S and localises to a different subset of dopaminergic neurons from Rab10. We propose that variations in *Rab* expression may contribute to differences in the rate of neurodegeneration seen in different dopaminergic nuclei in Parkinson’s.

**Significance Statement:** A key question in Parkinson’s is why dopamine neurons die particularly fast in some parts of the *substantia nigra*. We focused on the commonest Parkinson’s-related mutation, LRRK2-G2019S. *In vitro* assays suggested that neurodegeneration may start by LRRK2-G2019S increasing phosphorylation of Rab10. We found Rab10 in fly dopamine neurons in visual and motor pathways, but not in the sleep system. Rab10 knock-out rescues G2019S-induced visual and movement degeneration, leaving sleep dysfunction unaffected. Thus, LRRK2 activates at least two pathways, one Rab10-dependent, leading to neurodegeneration *in vivo*. Rab3 is found in a different subset of dopaminergic neurons and shows no synergy with *LRRK2-G2019S*. We propose that variations in *Rab* expression contribute to differences in neurodegeneration seen in Parkinson’s.

---

Inherited mutations in the LRRK2 (Leucine-rich-repeat kinase2) protein are the most common known cause of Parkinson’s. A single amino-acid change, *G2019S*, increases LRRK2 kinase activity (1), leading to a toxic cascade that kills dopaminergic neurons, but the main steps in this pathological signalling pathway remain to be determined. A number of studies highlighted a diverse range of >30 proteins that might be phosphorylated by LRRK2, suggesting it is a generalised kinase (2). However, several research teams have recently reported that LRRK2 is a more specific kinase, phosphorylating Rab GTPases (3–8). However, it is not clear which of the more than 60 Rabs are actually phosphorylated *in vivo*. Here, we use a *Drosophila* model to address the link from LRRK2 to Rabs physiologically in dopaminergic neurons, as these are lost in Parkinson’s.

Rabs are a plausible LRRK2 substrate leading to neurodegeneration, as they act as molecular switches interacting with a range of proteins (GEFs, GAPs and GDIs) regulating supply and delivery of cargo to membranes. Indeed many of the 66 Rabs in humans have been linked to neurodegenerative disorders (9). Mutations in Rabs 29 and 39 cause Parkinson’s (10–12). Biochemically, about 8 seem to be directly phosphorylated by LRRK2 [Rabs 3, 5, 8, 10, 12, 29, 35 and 43 (3)]. In mammals, analysis of the role of the Rabs is complex because individual Rabs may have similar, or even compensatory functions, which may differ by tissue (8, 13). The situation is simpler in the fly, because there are fewer Rabs - only 23 mammalian orthologs.

Furthermore, flies provide the well-developed GAL4-UAS system for neuron-specific gene manipulation, with the *Tyrosine Hydroxylase* (*TH*) GAL4 specific for dopaminergic neurons. This permits us to analyse the effects of *Rab10* manipulation in three systems: vision, a reaching movement task (proboscis extension) and sleep. We selected these systems as they are dopaminergic in both flies and mammals, and because the physiological and degenerative consequences of *LRRK2* manipulations can be accurately quantified using robust, highly sensitive assays (14–16).

For vision, we use the Steady State Visual Evoked Potential (SSVEP) assay based on that developed to analyse visual problems in humans. SSVEP measures the output of the photoreceptors and second-order lamina neuron. It provides a much more sensitive assay than conventional flash electroretinograms with less impact from movement or electrical interference. We found that a number of Parkinson’s-mimic flies showed increased activity in the lamina neurons at 1 day old, including those expressing *LRRK2*-*G2019S* in their dopaminergic neurons using the *Tyrosine Hydroxylase* GAL4 (*THG2* flies) (15, 17). Old *THG2* flies show loss of visual response, which is accelerated by a mild visual stress, achieved by keeping the flies in a ‘disco-chamber’ where the light is turned off every ∼2 s (18). Our proboscis extension assay was developed because a single dopaminergic neuron, TH-VUM, modulates the extension reflex (19) and we found that *THG2* flies show bradykinesia and tremor (14). Sleep – defined as a 5-minute period of inactivity – is routinely monitored by the Drosophila activity monitor, and *THG2* flies show aberrant sleep in 12:12 Light-Dark (LD) cycles.

Using these tools we tested if the LRRK2 ⇿ Rab pathway(s) reported in tissue culture also occur *in vivo*, focusing on functional outputs of dopaminergic neurons. If so, is LRRK2 specific to one or two Rabs, or is LRRK2 a more generalized kinase? Is any one Rab essential?

We determined that Rab10 has the strongest synergy with LRRK2-G2019S, Rab3 the weakest, and that each cluster of dopaminergic neurons expresses a different subset of Rabs. Dopaminergic Rab10 is essential for *THG2* visual and reaching phenotypes, but sleep phenotypes are independent of Rab10. This is correlated to high levels of Rab10 in the dopaminergic neurons associated with vision and proboscis extension, whereas Rab10 was not detected in dopaminergic neurons linked to sleep.

## Results

### A Visual Expression screen identifies that Rab10 has the strongest genetic interaction with LRRK2; Rab3 the weakest

In young flies where *Tyrosine Hydroxylase* GAL4 is used to express *UAS-LRRK2-G2019S* (*THG2*) the visual response in lamina neurons is much increased (15). This led us to screen flies with each *Rab* ± *G2019S* and record their visual response by SSVEP.

We found that dopaminergic expression of most *Rabs* (*TH* > *Rab*) phenocopies *THG2* (Fig. S1G), with a strong increase in visual response. This is true for both the photoreceptors (mean response 1.95 x *THG2*), and the lamina neurons (11.0 x *THG2*). For most Rabs, co-expression with *LRRK2-G2019S* (*THG2* > *Rab*) further enhances the visual response (photoreceptor 2.8 x; lamina neuron 14.5 x *THG2*; Fig. S1G, Table S1).

We next test if any Rab disrupts the neuronal feedback circuit between photoreceptor and lamina neurons. Normally, as the photoreceptor response increases, so does the lamina neuron response. This relationship is apparent throughout our screen (Fig. 1A), being remarkably similar in young (day 1) and older (day 7) flies. However, there is a one marked outlier, *Rab10*, where the lamina neuron response at day 1 is ∼5 times the value expected from the regression, and at day 7 is substantially below the line. Thus, in young *THG2* > *Rab10* flies there is much greater neuronal activity than expected, but in 1-week old *THG2* > *Rab10* flies we observe reduced activity, suggesting neurodegeneration has begun.

**Fig. 1.**
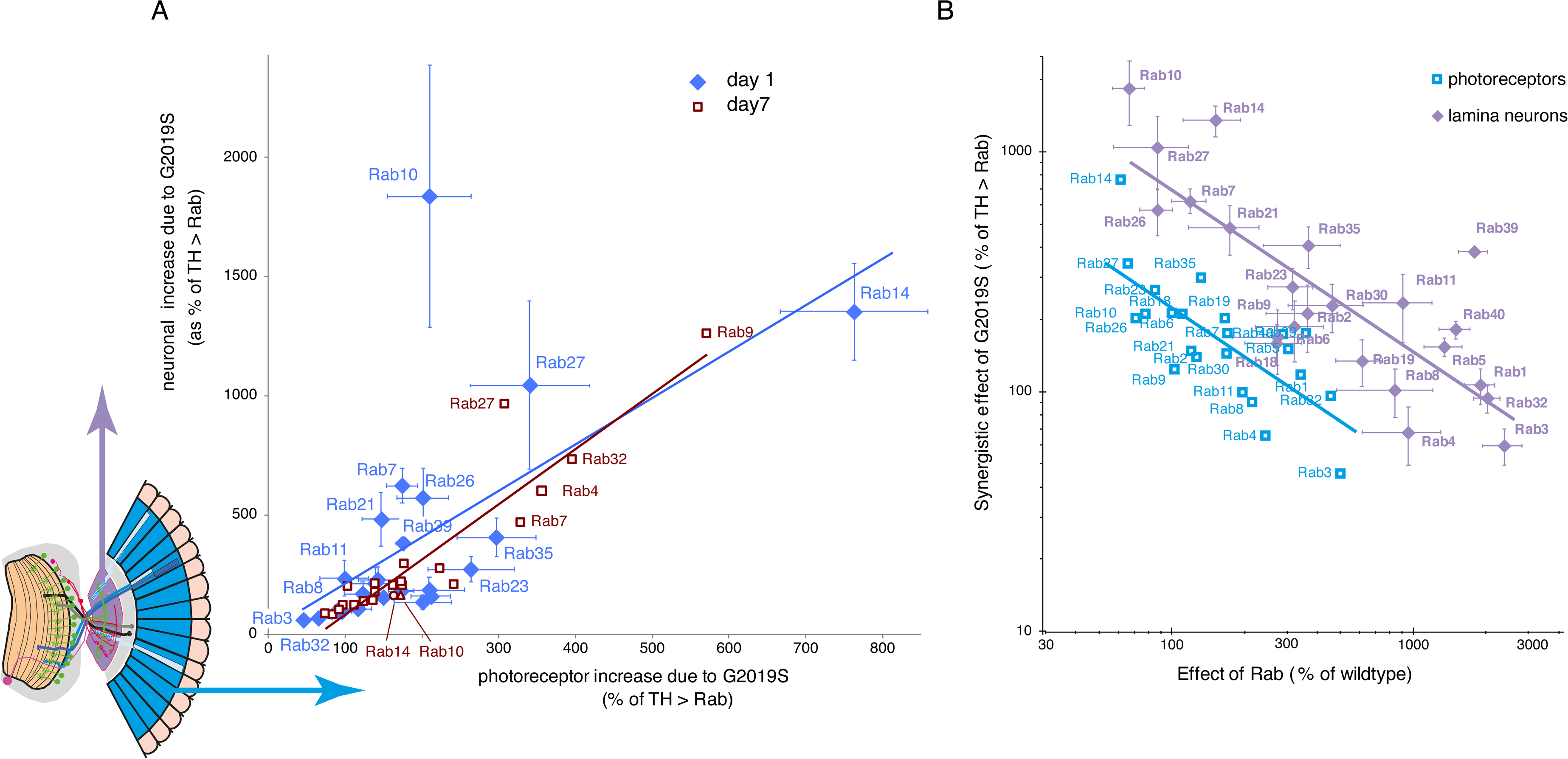
Expression screen highlights *Rab10* with the strongest synergy with *LRRK2-G2019S*, and *Rab3* as the weakest. Each *Rab* was expressed in dopaminergic neurons (using *TH*-GAL4) by itself (*TH* > *Rab*), or along with *G2019S* (*THG2* > *Rab*) and the visual response measured after 1 day or 7 days in the dark. A. **Standout role of *Rab10***. The response is split into two components, representing the photoreceptors and lamina neurons (inset blue and purple). The increase in lamina neuron response is highly correlated with the response of the photoreceptors, with one outlying exception, *Rab10* at 1 day. B. **Rab10 has the strongest synergy with *G2019S*.** Relationship of *Rab* and *G2019S* showing their inverse relationship. Rabs (3, 32, 1) which have a big effect on vision when expressed on their own have little further consequence when *G2019S* is also expressed; but other Rabs (10, 27, 14, 26) which have little visual impact on their own have a strong synergy with *G2019S*. Details in Figs. S1, S2 and Table S1.

Next, we tested if there is genetic interaction between *Rabs* and *LRRK2-G2019S.* For some Rabs (*10, 14, 27, 26*) expression of both *G2019S* and the *Rab* in dopaminergic neurons leads to a big increase in the lamina neuron response (Fig. 1B). Interestingly, these Rabs have little effect when expressed alone. The converse is also true: for the *Rabs* with the biggest effect (*3, 32, 1*), adding *G2019S* has no further effect.

We wanted to examine which factors controlled this synergy. A number of hypotheses have been put forward in the LRRK2 /Rab literature. First, LRRK2-G2019S is thought to phosphorylate only some Rabs *in vitro*, possibly due to a preference for Thr over Ser (20). Neither *in vitro* evidence for phosphorylation of the Rab by LRRK2, nor the amino-acid at the active site affects the regression (Fig. S2A,B). Nor is the relationship affected by the phylogenetic similarity of the Rabs (Fig. S2C). However, Rabs previously linked to Parkinson’s (21) have a stronger response to *G2019S* than others (Fig. S2D). Indeed, the Rab furthest above the regression line is one that causes Parkinson’s, *Rab39* (10).

We now ask if the synergy seen with joint expression of *Rab10* and *G2019S* could arise from phosphorylation of fly Rab10 by *LRRK2*-*G2019S*. The sequence of human and fly Rab10 is highly conserved, both where LRRK2 and a phospho-antibody to hRab10 bind, so that the antibody recognises fly pRab10 (Fig. S3). When *G2019S* is expressed pan-neuronally, a stronger pRab10 signal is seen than when a kinase dead form of *LRRK2* (*K1906M-G2019S*) is expressed. Thus, we conclude that *Rab10* shows the strongest synergy with *LRRK2-G2019S* in our visual electroretinogram assay, most likely due to direct phosphorylation by LRRK2. Contrastingly, Rab3 expression exhibits no synergistic interactions. This is an important difference, because both Rab10 and Rab3 are reported to be phosphorylated by LRRK2 *in vitro*.

### A novel Rab10 null

Since *Rab10* was the top hit in our overexpression assay, we wanted to test if reduction of *Rab10* affected retinal degeneration. We generated a *Rab10* null by deleting the coding region from just after the start of the GTPase domain to the end of the protein using CRISPR / Cas9 (Fig. S3). The deletion includes the LRRK2 binding site. The null mutant (*Rab10*) is viable, unlike that of the mouse (22).

We tested for a *Rab10* phenotype in visual assays. When kept in the dark at 22 °C, *Rab10* flies had a normal flash electroretinogram (Fig. S4). However, flies kept in the dark at 29 ° and tested in the more sensitive SSVEP assay have a larger neuronal response than their matched control (*Rab10; TH* and *TH/+*, Fig. S5); those *Rab10; TH* flies kept in the disco-chamber to accelerate neurodegeneration have much worse visual response than controls (Fig. 2D). Dopaminergic expression of *Rab10 RNAi* does not affect the vision of flies kept in the dark (Fig. S5) but does accelerate the loss of lamina neuron function in the disco chambers (Fig. 2D, compare *TH; Rab10 RNAi* with *TH/+*). These experiments suggest dopaminergic Rab10 is important for maintenance of visual response to light in aged flies.

**Fig. 2.**
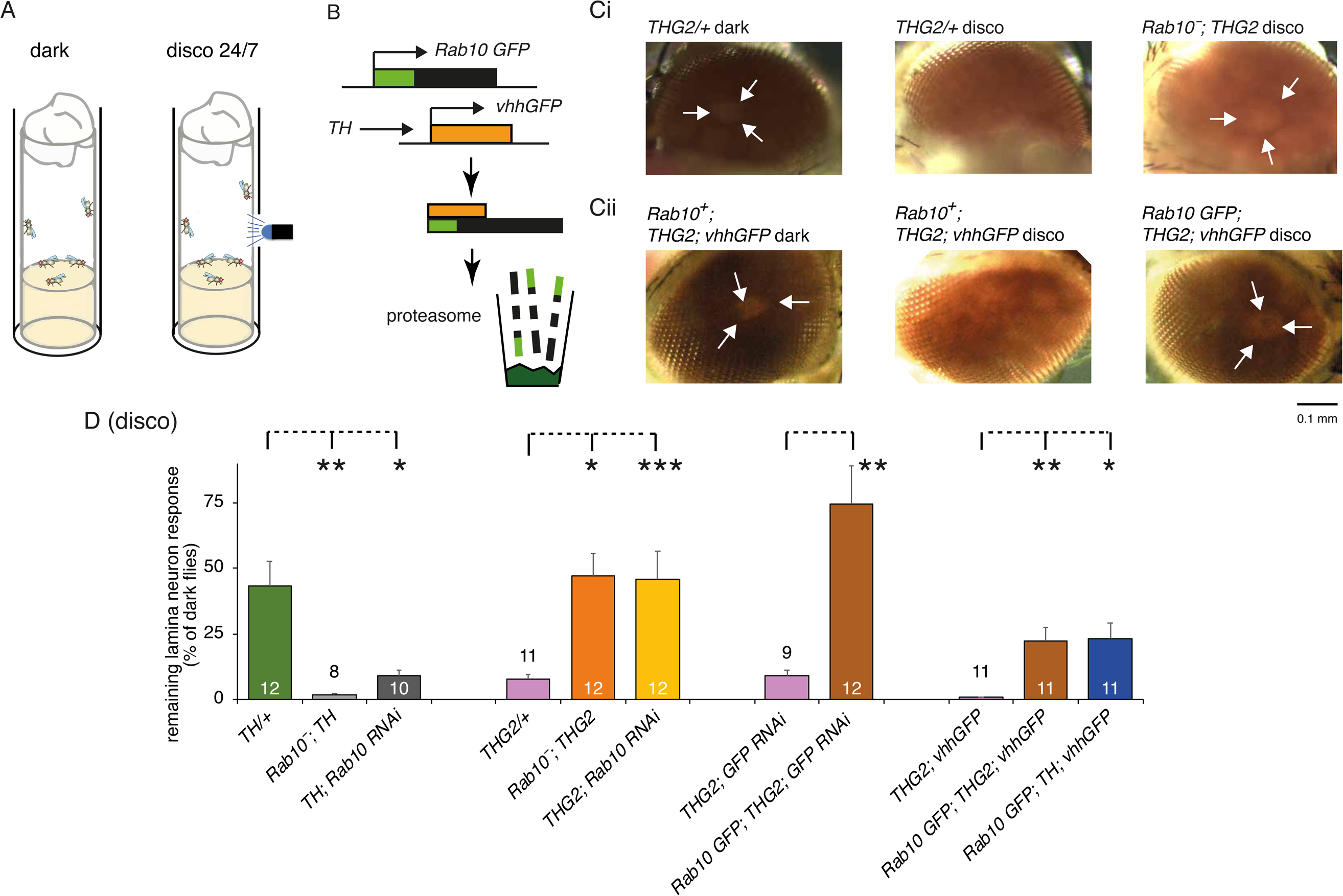
Accelerated visual decline due to *LRRK2-G2019S* is rescued by dopaminergic knock-down of *Rab10*. A. Accelerated visual decline was achieved by keeping flies in disco-chambers (18) compared with those in the dark. B. Destruction of Rab10 GFP protein in the deGradFP technique by dopaminergic expression (*TH*-GAL4) of a nanobody targeted at GFP (*vhhGFP*), which sends Rab10 GFP protein to the proteasome. C. **Anatomically, dopaminergic *Rab10* knock-down rescues the loss of eye structure.** All flies kept in the dark for 10 days have strongly pigmented eyes, with a marked DPP (Deep Pseudo-Pupil (23)). This shows that the internal, reflective structure of the eye is intact. Flies with dopaminergic expression of *LRRK2-G2019S* (*THG2*) kept in disco chambers lose their pigmentation (indicative of lysosomal dysfunction) and have no DPP. This is rescued in the *Rab10* null (Ci: *Rab10*; *THG2*), or by knockdown of Rab10 protein in dopaminergic neurons using the deGradFP technique (Cii: *Rab10 GFP; THG2*; *vhhGFP* flies). D. **Physiologically, dopaminergic *Rab10* knock-down rescues the neuronal vision.** *THG2* flies in disco chambers lose their physiological response after 7 days (magenta bars). This is rescued by global knock-out of Rab10, (*Rab10*; *THG2* orange bar) or by dopaminergic knock-down of Rab10 by *RNAi* (*THG2*; *Rab10RNAi* yellow bar). Rescue is confirmed using flies in which Rab10 had been replaced with *Rab10 GFP* using RNAi targeted at Rab10-GFP mRNA or using deGradFP mediated knockdown of Rab10 protein (*Rab10 GFP; THG2; GFP* or *Rab10 GFP; THG2; vhhGFP* brown bars). Control experiments show that Rab10 knock-out (*Rab10; TH*) or dopaminergic knock-down (*TH* > *Rab10 RNAi*) accelerates the loss of visual function. This is particularly marked for the *Rab10; TH* flies, as these have a stronger visual response than the other genotypes when kept in the dark. The lamina neuron response of dark-reared flies and the neuron / photoreceptor relationship is in Fig. S5, with exact genotypes in Table S2.

### Global or dopaminergic reduction in Rab10 rescues G2019S-induced anatomical deficits

The key question is how Rab10 reduction affects *G2019S*-induced neurodegeneration. We started by measuring retinal structure anatomically, using the ‘Deep-Pseudo-Pupil’ (DPP) assay, comparing flies raised in the dark those kept in a disco-chamber to accelerate neurodegeneration (Fig 2A). Dark-reared flies show a DPP, since the normal retinal structure focuses light from below towards the observer through 7 ommatidia (23), (Fig. 2Ci). Disco flies expressing *LRRK2-G2019S* in the dopaminergic neurons (*THG2*) lose pigment and their DPP by 10 days, with disorganized patterns of light and dark across the surface of the eye (Fig. 2Ci). The lack of DPP in disco-reared *THG2* flies indicates that light is scattered randomly within the eye since the internal retinal structure is lost. However, in the *Rab10 null* (*Rab10; THG2*), the DPP is preserved, showing *Rab10* knock-out rescues the *G2019S-*induced loss of retinal structure.

As global Rab10 knock-out protects the structure of the eye from *G2019S*-induced degeneration, we next asked if dopaminergic reduction of Rab10 is sufficient to rescue the structural deficit. We used the deGradFP technique (Fig. 2B) to achieve dopamine-specific Rab10 knock-down. For this, we took flies in which GFP has been inserted at the start of the *Rab10* gene (*Rab10 GFP*) (24). We depleted Rab10 GFP protein specifically in dopaminergic neurons by targeted expression of a GFP-targeted nanobody (*vhhGFP*). This marks GFP-proteins for proteolysis (25, 26). Flies with Rab10 GFP, nanobodies and G2019S show normal pigmentation and DPP whether kept in the dark or in a disco-chamber. This shows that depletion of Rab10 protein is sufficient to protect the flies from *G2019S*-induced degeneration. As a control, flies with the nanobody and G2019S, but with wild-type Rab10, lose pigmentation and DPP, as the nanobody does not bind to the normal Rab10.

These experiments suggest that knock-down of Rab10 in dopaminergic neurons plays a role in protecting against the *LRRK2*-*G2019S* induced neurodegeneration.

### Global or dopaminergic reduction in Rab10 rescues G2019S-induced visual function

To consolidate the anatomical results, we tested the electrophysiological response of the eyes. All dark-reared flies show a strong lamina neuron signal – at least >50% of the control (Fig. S5). The response is less in flies with multiple transgenes due to their darker eyes. Disco-reared flies show strong effects of genotype: the *THG2* genotypes in three control backgrounds all have very poor vision, < 10 % of dark-reared progeny of the same cross (Fig. 2D, magenta bars). In the global *Rab10* null background, this response is fully rescued (Fig.2D compare green and red bars). Thus, both electrophysiological response and structural observation of the DPP indicate that *Rab10* null rescues the effect of dopaminergic *LRRK2-G2019S*.

Since global knock-out of Rab10 rescued the *G2019S* phenotype, we next asked if dopaminergic-neuron-specific knock-down of Rab10 is enough to rescue the visual deficits induced by *G2019S* in disco-chamber flies. The first strategy was to use a *Rab10* specific RNAi driven by *TH-*GAL4. In these flies (*THG2; Rab10 RNAi* Fig. 2D, yellow bar) the lamina neuron response is completely rescued. To confirm that Rab10 knock-down rescues *G2019S* mediated degeneration, we used the flies with *Rab10 GFP* and depleted the mRNA or protein using *GFP RNAi* or *vhhGFP* expression. These flies (*Rab10 GFP; THG2; GFP RNAi* or *Rab10 GFP; THG2; vhhGFP* Fig. 2D brown bars) show much larger lamina neuron visual responses than the corresponding controls with wild-type Rab10 where there is no GFP, and so the Rab10 is fully functional. Thus, all three strategies indicate that dopaminergic Rab10 is essential for the *G2019S*-induced loss of physiological response, as well as anatomical degeneration.

### Rab10 reduction ameliorates a movement LRRK2-G2019S phenotype

The role of Rab10 in a visual phenotype lead us to ask if Rab10 is essential for all *LRRK2-G2019S* phenotypes. Dopaminergic signalling is important in movement, including a reaching movement used to obtain food, the Proboscis Extension Response (PER, Fig. 3A). Expression of *G2019S* in a single dopaminergic neuron, TH-VUM (19) results in movement deficits – akinesia (loss of the PER, Proboscis Extension Response), slower response (bradykinesia) and tremor (14). When the PER is elicited by an optogenetic stimulus, or by sucrose application, dopaminergic knock-down of *Rab10* fully rescues the *G2019S*-induced akinesia (Fig. 3B, Fig.S6). Additionally, Rab10 reduction ameliorates bradykinesia, reverting the slow extension seen in *G2019S* flies (Fig. 3C). This was not due to a deficit in extending the proboscis as all genotypes extended their proboscis to the same extent (Fig. 3Ci). Dopaminergic expression of *Rab10* phenocopies *G2019S* expression, reducing the proportion of flies that respond (Fig. S6).

**Fig. 3.**
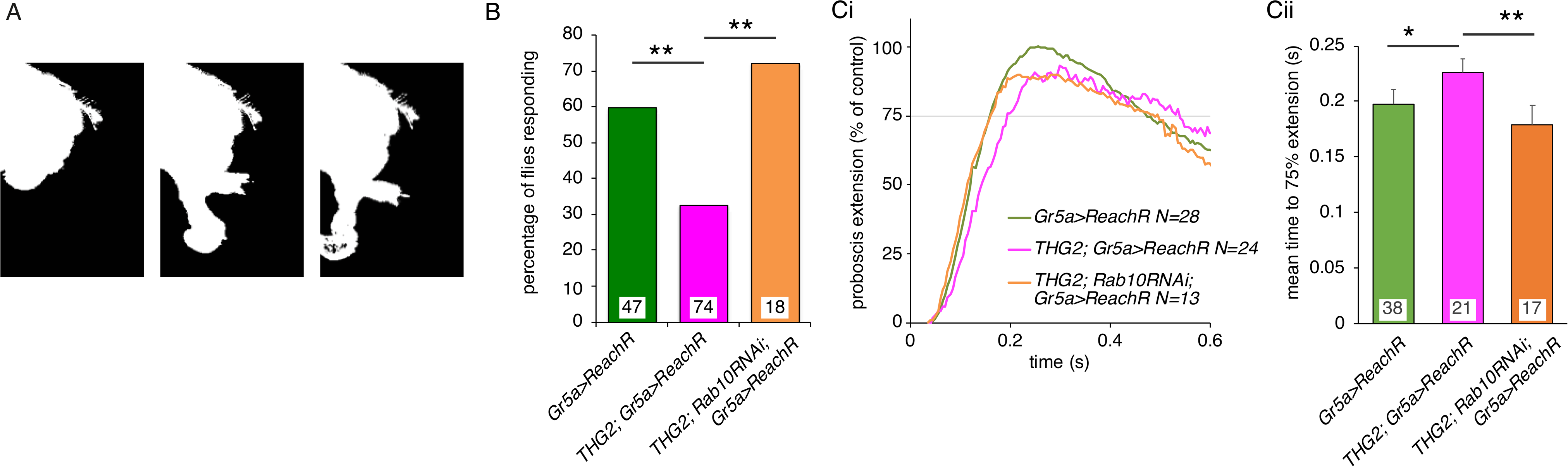
*Rab10* knock-down rescues proboscis reaching deficits induced by dopaminergic expression of *LRRK2-G2019S*. A. To reach for food or liquid, the fly extends its proboscis in response to a stimulus to sensory neurons on the legs. B. Dopaminergic reduction in *Rab10* rescues the akinesia (less frequent response) of flies expressing *G2019S* in their dopaminergic neurons. C. B. Dopaminergic reduction in *Rab10* rescues the bradykinesia (slower response) of flies expressing *G2019S* in their dopaminergic neurons. Ci. Raw traces; Cii. mean data. To respond to the optogenetic stimuli all flies use LexA / Op *Gr5a*>*ReachR.* Exact genotypes in Table S3.

These experiments show that Rab10 reduction ameliorates movement, as well as visual deficits induced by *LRRK2*-*G2019S*.

### Rab10 reduction fails to ameliorate a ‘sleep’ LRRK2-G2019S phenotype

Dopaminergic neurons also influence circadian rhythms and ‘sleep /wake’ patterns (27, 28). At least three clusters of dopaminergic cells (PAM, PPL1, PPM3) contribute to this phenotype, though the exact pathways have yet to be fully determined. In 12:12 light on:off cycles, *THG2* flies show a marked sleep deficit (Fig.4). In the dark, they spend less than 50% of the time ‘asleep’ than control flies do, but in the light they sleep more. Manipulation of *Rab10* has no effect on sleep (Fig. 4) but gives subtle changes with the lights off responses of these flies (Fig. S7A). Dopamine level has little effect on circadian period (29) and neither *LRRK2-G2019S* nor *Rab10* manipulations affects the circadian period of flies in constant darkness (Fig. S7B).

**Fig 4.**
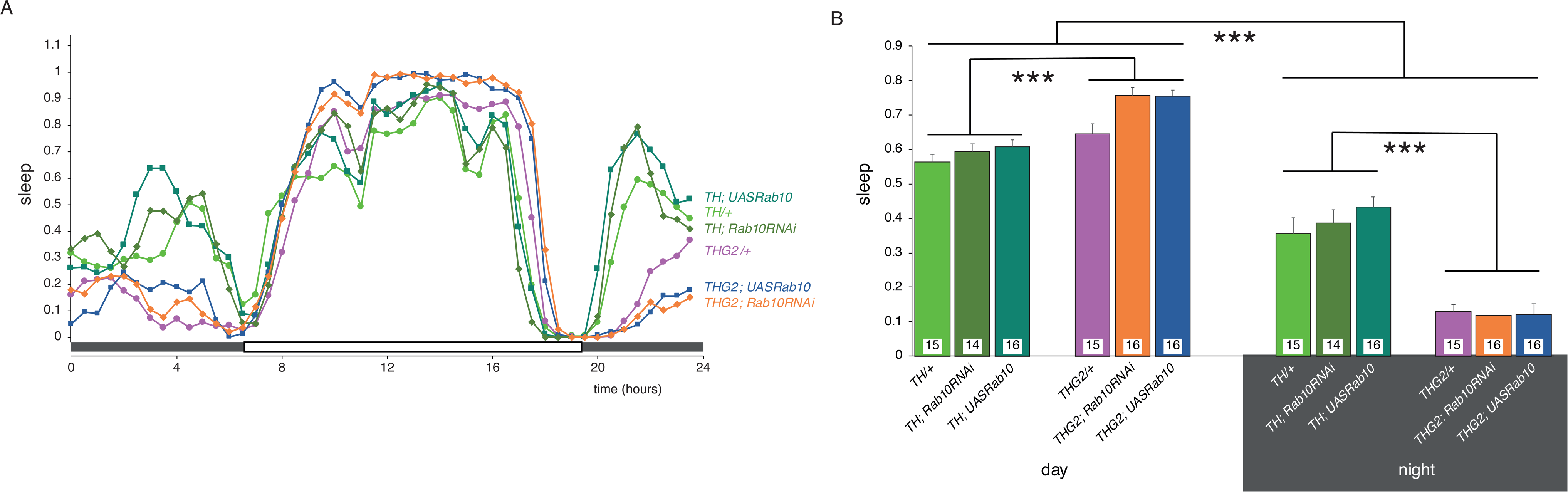
*Rab10* knock-down does not rescue sleep deficits induced by dopaminergic expression of *LRRK2-G2019S*. A. *THG2* flies show increased sleep during the day, and reduced sleep at night. Neither reduced nor increased expression of Rab10 affects the daily pattern (Ai), with summary data in Aii. Sleep is defined as periods of inactivity longer than 5 min. Actograms from these flies are shown in Fig. S5. Exact genotypes in Table S4.

Thus, we find that *Rab10* reduction does not rescue the sleep phenotype, unlike the visual and proboscis extension phenotypes.

### Rab10 is only found in a subset of dopaminergic neurons

To examine the overall distribution of Rab10, we used *Rab10*-GAL4 (30) to express a nuclear RFP, and α-TH antibody to mark the dopaminergic neurons. We found that many, but not all dopaminergic neurons were Rab10 positive (Fig. 5A). Rab10 positive neurons include the small MC neurons in the optic lobes (Fig. 5Aii), the TH-VUM neuron (that controls the PER) and DADN neurons (control leg movements) (Fig. 5Aiii). Coincidence was also observed in some PAL, PPM3, PPL2, and T1 neurons. However, other neurons, notably those in the PPL1, PPM2 and PAM clusters did not show *Rab10* driven RFP. The PPL1 and PAM neurons have been linked to sleep / wake patterns (27, 28). To confirm this data, we adopted an intersectional strategy, based on split GFP, whereby dopaminergic *TH*-LexA and *Rab10*-GAL4 were used to deliver the two halves of GFP. Only neurons which are both dopaminergic and Rab10+ will generate GFP and fluoresce. This delivered the same pattern of coincidence (Fig. 3Aiv). A different subset of dopaminergic neurons is Rab3 positive (Fig. S8).

**Fig. 5.**
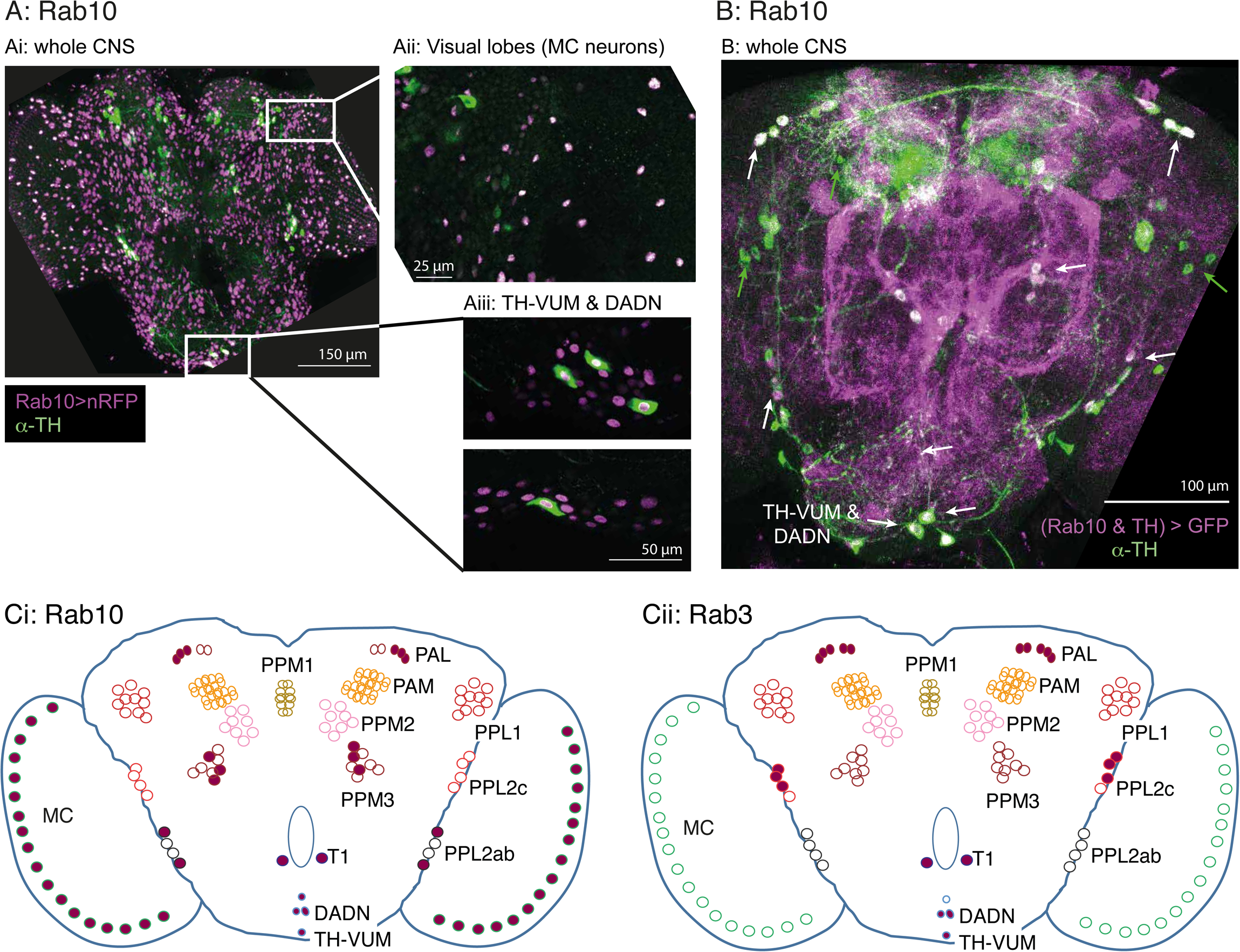
Rab10 and Rab3 are located in different subsets of the dopaminergic neurons. A. Not all dopaminergic neurons, identified by a cytosolic α-tyrosine hydroxylase antibody (α-TH, green), are indicated by *Rab10*-GAL4 expression of nuclear RFP (magenta). Rab10 is detected in the dopaminergic MC neurons (in the optic lobe) (Aii) and in the TH-VUM (controls proboscis extension response (19)) and DADN neurons (control leg movements (64)) (Aiii). B. An intersectional strategy to express *Rab10*-GAL4 (magenta) confirms that not all the dopaminergic neurons (green) are positive for Rab10. White arrows indicate cells marked with GFP and α-TH, including the TH-VUM and DADN neurons; green arrows cells which are dopaminergic but not Rab10. C. Summary of the dopaminergic neurons coincident with (i) Rab10 and (ii) Rab3. Diagram and nomenclature after (65, 66). Antisera and exact genotypes in Table S5.

We next asked if Rab10 was indeed phosphorylated in any dopaminergic neurons. In the fly brain, the phospho-Rab10 antibody marks 3 clusters of neurons along the dorsal edge of the brain: a lateral pair and a medial cluster (Fig. S8A,B, white arrows). Also stained are a pair or ventral neurons, close (but not identical) to the dopaminergic TH-VUM and DADN (Fig. S8A,B, yellow arrows). None of these phospho-Rab10 cells were marked by the α-TH antibody used to identify dopaminergic neurons. No cells were strongly marked in the *Rab10* flies (Fig. S8C, one cell was weakly stained, grey arrow), showing the specificity of the antibody.

Thus, we conclude that some dopaminergic neurons natively express Rab10, notably those concerned with vision and Proboscis Extension - while in others the level of Rab10 is too low to be detected – those controlling ‘sleep’. Only a few of the Rab10+ neurons show high levels of phospho-Rab10.

## Discussion

Our key observation is that knock-out of Rab10 *in vivo* fully rescues the reduced responses induced by dopaminergic *LRRK2-G2019S* in visual and motor (reaching, proboscis extension) assays, but the sleep phenotype is unaffected. Rab10 is expressed in dopaminergic neurons controlling vision (MC) and proboscis movement (TH-VUM), but undetectable in those controlling sleep, so that the anatomical and physiological patterns of Rab10 are related. Our results support the idea that LRRK2 phosphorylates separate targets in distinct neurons.

The existence of Rab10 in the tyrosine hydroxylase positive neurons controlling vision and proboscis movement argues that LRRK2 might indeed phosphorylate Rab10 directly. Thus our *in vivo* results both support the *in vitro* (biochemical and cell culture) data in which LRRK2 directly phosphorylates Rab10 (3–8), but also suggest a separate pathway for *LRRK2*-*G2019S* action. Further, while Rab3 is also phosphorylated by LRRK2 *in vitro* (3), it did not synergise with *LRRK2*-*G2019S* in the visual assay, and was not detected in the MC neurons.

In the visual system, we confirmed dopaminergic rescue of both anatomical and physiological deficits using three different genetic strategies. In the akinesia movement assay, dopaminergic *Rab10* knock-out (two strategies) rescues the *G2019S* deficit. In both systems, rescue is back to wild-type levels, identifying Rab10 as the key component. However, we have noted that in human peripheral mononuclear blood cells, LRRK2 phosphorylates both Rab10 and Rab12 (6). Since the fly has no Rab12, with Rab10 being the closest homolog, we cannot exclude a G2019S □ Rab12 relationship in mammalian dopaminergic neurons.

The effects of LRRK2-G2019S and Rab10 on vision and proboscis extension could be accounted for by a reduction in dopamine release. Dopamine inhibits photoreceptors (31) and excites proboscis extension (19). This fits with the observation, in worm, that Rab10 is required for release of transmitter from Dense Core Vesicles (DCVs) (32) and we note that dopamine is packaged in DCVs. In mammals, Rabs 10, 14 facilitate delivery of GLUT4 vesicles to the plasma membrane in adipocytes (13). However, a developmental effect cannot be excluded as Rab10 modulates neuronal growth (34, 35).

### Expression screen highlights synergy between Rab10 and G2019S

A wider view of possible LRRK2 ⇿ Rab pathways is provided by the visual screen. All Rabs, either by themselves or when co-expressed with *G2019S*, increase the visual response. One possible physiological mechanism is that Rab overexpression reduces retinal dopamine release onto the photoreceptors. This will increase the amplitude and speed of the photoreceptor response (31). Dopamine may also affect the lamina neurons, and third order MC cells, but this remains to be determined. It is also possible that the Rabs modulate the release of co-transmitters or growth factors from dopaminergic neurons.

Our screen placed the Rabs along a spectrum, ranging from those with a strong synergy with *G2019S* to those which had a strong effect when expressed by themselves.

Rabs 10, 14 and 27 have the strongest synergy with *G2019S*, though by themselves they have little effect on visual sensitivity. These three Rabs are best known for their roles in exocytosis. However, another function for Rab10 is suggested by the stand-out, large lamina neuron: photoreceptor response ratio when *G2019S* and *Rab10* are expressed together. This might the role of Rab10 in endocytosis (36–39). Here, the effects of Rab10 are mediated through the EHBP1-EHD2 complex. In the follicle cells of *Drosophila, ehbp1* expression and knockdown phenocopy Rab10 manipulations (40), while EHBP1 was also identified by a systematic proteomic analysis as indirectly phosphorylated by LRRK2 in HEK293 cells (3) and a lysosomal assay (41).

Among the Rabs which have strong visual effect when expressed in dopaminergic neurons, but show little synergy with *G2019S* are 1, 3, 5, 6 and 11. Rab3 is notable for its role in synaptic transmission, while the other Rabs here play roles in the fly optic system in intracellular cycling [see for review (42)]. Indeed, Rab3 was at the opposite end of the Rab spectrum from Rab10 – this is notable because *in vitro* mammalian cell assays have highlighted similar roles of Rabs 3 and 10 in lysosome exocytosis, (43, 44), as well the observation that both phosphorylated in HEK cells (3). On explanation for the discrepancy between the *in vivo* and *in vitro* assays is that Rab3 is localised to synapses, possibly far from LRRK2 at the trans-Golgi network (7).

Our observation that every Rab seems to have some effect on dopaminergic signalling in the visual system goes some way to explain why many studies of individual Rabs have demonstrated effects with LRRK2; Rab3a (45); Rab5 (46); Rab7 (47); Rab29 (11). Although cellular studies support an binding of Rab29 to LRRK2 (48), the closest fly homolog (Rab32) shows little synergy with *G2019S* in our screen.

The availability of Rab transgenic flies facilitates screening in *Drosophila*. Screens have identified key roles for Rab2 in muscle T-tubule development (49); Rabs 2, 7, 19 in loss of huntingtin (50), 1, 5, 7, 11 and 35 in the *Drosophila* renal system (51), Rab32 in lipid storage (52) and Rab39 in tracheal formation (53). The varied outcomes of these screens indicate the validity of the LRRK2-G2019S ⇿ Rab10 relationship reported here.

### Each dopaminergic neuron has its own palette of Rab expression

Finally, we note that not all dopaminergic neurons are equally susceptible in Parkinson’s. A long-standing observation is that the dopaminergic neurons in the VTA (ventral tegmental area) do not degenerate in the same way as those in the *substantia nigra*. More particularly, even within the *substantia nigra* there is a range of outcomes, with dopaminergic neurons in the *pars compacta* dying more than those in the dorsal and lateral zones (54). The same is true for fly neurons: the neurons in the PPM1 and PPM2 degenerate more than those in other clusters (though no data is available for the MC and TH-VUM neurons which are our focus). Previously, faster neurodegeneration has been ascribed to increased cytosolic dopamine levels (55), to intracellular effects of glutamate (56), to increased calcium influx (57), through more action potentials (58), or to longer axons with more synapses (59). It has not escaped our notice that faster degeneration in some neurons may be the result of their different palettes of Rab proteins.

## Materials and methods

### Flies

(*Drosophila melanogaster*) were raised and manipulated according to standard fly techniques. Data in Figs. 1 and S1 is from female flies; all other data from male flies.

A new *Rab10* was generated by CRISPR-mediated mutagenesis by WellGenetics Inc. using modified methods by (60). A 1644-bp region, spanning from +57 nt relative to ATG of Rab10 to +70 nt relative to the 1st nt of Stop of Rab10, was replaced by a STOP-RFP knock-in cassette, deleting most of the coding region.

Briefly, a pair of gRNAs (upstream: CTGATCGGTGATTCAGGAGT, downstream: GAACGGGGCGTGGTTTGGCC) was cloned into a U6 promotor plasmid. For repair, a STOP-RFP cassette containing 3xSTOP codon and 3xP3-RFP and 1-kb homology arms were cloned into a pUC57-Kan donor plasmid. Wild-type w1118 embryos were microinjected with Rab10-targeting gRNAs and hs-Cas9 supplied in DNA plasmids, together with the donor plasmid. Flies carrying the positive selection marker 3xP3-RFP were validated by genomic PCR and sequencing. For the upstream PCR, primers were designed at 5’ outside of upstream homology arm (Fwd primer 5’-GATGTTCAGGCGATTCTGGT) and at 3xP3 promotor (Rev primer 5’-CGAGGGTTCGAAATCGATAA). For the downstream PCR primers were designed at alpha-tubulin 3’ UTR, which is used for STOP-RFP transcript stability (Fwd primer 5’-AACGCAAGCAAATGTGTCAG) and at 3’ outside of downstream homology arm (Rev primer 5’-CGATGATTGTTTGCCATTTG).

Fly stocks are listed in Table S6.

### Visual assays

On the day of emergence, flies were placed in the dark or in disco-chambers at 29 °C. 1 day or 1 week old flies were prepared for SSVEP (Steady State Visual Evoked Potential) measurements as described (Fig. S1) (15). The same protocol was used, except that stimuli were generated, and responses recorded by an Arduino Due system instead of a PC. Data was analysed in Matlab and R. Full code at https://github.com/wadelab/flyCode. DPP (Deep-Pseudo-Pupil) assays were performed on a Zeiss Stemi microscope with AxioCamERc5s camera.

### Western Blots

were made using standard protocol and antibodies: α-phospho-Rab10 (Abcam: ab230261, 1:1000) with α-synaptotagmin 91 as a loading control (61). Control Rab10 samples were derived from female Wistar rats (Charles River UK), maintained in accordance with the UK Animals (Scientific Procedures) Act (1986). Animals were euthanised via exposure to CO_2_ in a rising concentration, followed by cervical dislocation in accordance with Home Office guidelines and with approval of the Biology Ethics Committee (University of York). The brain was dissected into PBS and 10 mg of tissue lysed for the blots (62).

### Immunocytochemistry

was performed as described recently (14). Tyrosine hydroxylase was detected with Mouse anti TH Immunostar (22941, 1:1000), and phospho-Rab10 (Abcam: ab230261, 1:250).

### Akinesia

was recorded from 1 week old flies, kept in the dark at 29 °C. Flies were restrained as described (14) and starved at 25 °C for 3 hours before being offered a droplet of 100 mM sucrose three times. Each response was scored Yes /No and the median response for each sample used. χ^2^ with post-hoc tests used the R package ‘fifer’.

### Bradykinesia

was assessed using an optogenetic stimulus. Flies were fed retinal (1 mM) pipetted onto the surface of their food for 1 week in the dark at 29 °C. They were restrained at 25 °C for 3 hours before the proboscis extension responses were observed with a Grasshopper 3 (Point Grey) camera mounted on a Zeiss Stemi microscope at 200 frames / second. A single flash was delivered from a ThorM470L3 LED, driven at 8 V for 7 ms. The stimulus was transcoded by LexOpReachR expressed in the Gr5a neurons. The area of the video occupied by the proboscis was automatically analysed by python code https://github.com/biol75/PER, and the statistics in R.

### Circadian activity

patterns were recorded with a TriKinetics DAM system, in dim light at 29 °C and analysed using the ShinyR-DAM code (63) using the branch at https://karolcichewicz.shinyapps.io/ShinyR-DAM_3_1_Beta/.

### Statistics

were calculated in R, with the mean ± SE reported by error bars or median ± interquartile range in box plots. Post-hoc tests were calculated for χ^2^ using the fifer package and for ANOVA, the Dunnett test.

## Acknowledgements

We are grateful for the gifts of flies from Kristin Scott, Wanli Smith, Cheng-Ting Chien, Julie Simpson, Robert Kittel, Serge Birman, Yoshi Aso, Mattias Landgraf and Stefan Heidmann. We also thank the York Biology Technology Facility, Bloomington Drosophila Supply Center and Flybase for their provision. Olivia Compton and Martin France helped considerably with pilot studies. Sean Sweeney, Chris MacDonald and Amy Cording kindly read a draft manuscript. We are particularly grateful to Parkinson’s UK and to their volunteers for support (K-1704), and grants from the Deutsche Forschungsgemeinschaft (SFB /TRR186) to P.R.H..

