## Supplementary Figures and tables for "Neurodegeneration caused by LRRK2-G2019S requires Rab10 in select dopaminergic neurons"

**Supplementary Figure Legends**

Fig. S1. **SSVEP (Steady State Visual Evoked Potential Analysis) determines the contrast response function of the insect eye**. A. The fly eye consists of ~800 ommatidia, each containing 8 photoreceptors. Their axons project to the lamina and medulla, where they synapse with the second- and third order neurons (lamina and medulla neurons). The medulla contains intrinsic dopaminergic neurons (MC), while some dopaminergic neurons from the CNS project to the lamina. B. Recording the fly visual response: A fly, restrained in a pipette tip, is illuminated with blue light from a LED, and the voltage across the eye is amplified and recorded. C. Repetitive stimuli given to the fly about a fixed mean light evoke a contrast response increasing with the peak-peak excursion of the stimulus waveform. D. The response to a series of identical stimuli is analysed by the Fast Fourier Transform, and averaged. This shows a response at the stimulus frequency (1F1) and additional components at multiples of the input, notably twice the input frequency (2F1). Genetic dissection shows that the 1F1 component is mostly generated by the photoreceptors and the 2F1 by the lamina neurons (15, 67). E. Plotting the amplitude of the 1F1 and 2F1 components against the stimulus contrast generates a CRF (Contrast Response Function), which differs from fly to fly. F. The averaged maximum CRF is dependent on genotype, with *THG2* (flies expressing *LRRK2-G2019S* in their dopaminergic neurons under the *Tyrosine Hydroxylase*-GAL4, *TH*) and *TH* > *Rab7* both showing a bigger response than control flies (*TH* crossed with wild-type, $w^{-}$). G. Overview of our screen, showing the photoreceptor (1F1) and lamina neuron (2F1) responses at 1 and 7 days for all the orthologs of mammalian Rabs. Data is calculated as the percentage of *THG2* flies. Data and number of flies is in Table S1.

Fig. S2. **An established role in Parkinson's is the only factor that influences the inverse relationship between *TH* > *Rab* and *THG2* > *Rab***. The LRRK2 ↔ Rab relationship found in Fig. 1B is replotted here to test if it is affected by factors that have been proposed to influence LRRK2 Rab function. A. Some

Rabs are phosphorylated by LRRK2 *in vitro* (3), but these Rabs are not more sensitive to *G2019S in vivo*. B. Rabs usually have a serine (S) or threonine (T) where they could be phosphorylated by LRRK2, though Rab40 has a histidine (68). Although a preference for LRRK2 to phosphorylate Rabs with a threonine was suggested by *in vitro* assays (20), *in vivo* there is no effect of the amino-acid. C. Sensitivity is not linked to the phylogenetic grouping (68) of the fly Rabs. D. Rabs previously linked to Parkinson's (21) have a stronger Rab ↔ G2019S interaction than those which do not influence Parkinson's. Summary data and exact genotypes in. Table S2.

Fig. S3. **High homology between *Drosophila* and human Rab10**. Ai. An antibody targeted at the phosphorylated switch II domain of human Rab10 recognizes both rat and fly Rab10, but no phospho-Rab10 protein is detected in the Rab10 null (*Rab10*¯) or when Rab10 RNAi is driven pan-neuronally (*nSyb* > *Rab10* RNAi). Pan-neuronal expression of *Rab10* (*nSyb* > *UASRab10*) increases the intensity of the phospho-Rab10 band. The α-drosophila-synaptotagmin antibody does not recognize any rat protein. Total Rab10 was not determined as the antibody binds to an unconserved region. Aii. Expressing *LRRK2-G2019S* pan-neuronally phosphorylates more Rab10 than expressing a kinase-dead form of *LRRK2* (*LRRK2-KD, LRRK2-G2019S-K1906M*). The wild-type was a *CS/w*¯ cross. B. Comparison of fly and human Rab10, showing conservation in the GTPase domain and prenylated region. Also shown are the *Rab10*¯ deletion, starting at position 21, the region targeted by the phospho-Rab10 antibody (amino-acids 66-80), the region targeted in *Rab10* RNAi and Thr73 which is phosphorylated *in vitro* by LRRK2.

Fig. S4. **Flash electroretinogram (ERG) recordings of *Rab10*¯ flies show no visual phenotype.** A. Representative flash ERG traces of newly hatched, 0 day old wild-type control and *Rab10*¯ flies. The response to a high intensity 1 s white light was recorded with a dc-coupled amplifier. B. Quantification of the flash ERG recordings. No visual decline occurs in newly hatched *Rab10*¯ flies (0 days) or in 4 days old flies, kept in the dark at 22 °C, when compared to wild-type control flies (*y w*). *N* = 25-30 flies per sample.

Fig. S5. A. **All flies kept in the dark have good visual responses**. Summary of lamina neuron response for each genotype shown in Fig. 2D. B. **Correlation of photoreceptor and lamina neuron responses persists in flies with *Rab* knock-down**. As in overexpression assays (Fig. 1B), a linear relationship exists between the photoreceptor and lamina neuron response. Summary data (including the exact genotypes) are shown in Table S2.

Fig. S6. **Rab10 reduction ameliorates the Proboscis Extension Response deficits induced by dopaminergic expression of *LRRK2-G2019S* (sucrose stimulation).** In response to application of sucrose to the legs, flies evert their proboscis, but the proportion of flies that respond is reduced in *THG2* flies (magenta bars). This is phenocopied by dopaminergic expression of *Rab10* (dark blue bars) and is ameliorated by either global or dopaminergic reduction of *Rab10* (orange bars). Exact genotypes Table S3.

Fig. S7. ***Rab10* knock-down does not rescue the minor circadian deficits induced by dopaminergic expression of *LRRK2-G2019S*.** A. Actograms recorded in 12:12 LD cycles show that dopaminergic expression of *G2019S* reduces overall activity and abolishes the extra activity seen with lights-on. Rab10 reduction abolishes the lights off peak in activity, providing a link to the electroretinogram data (Fig.2D). Despite these minor changes, the circadian timing persists. The sleep pattern of these flies is shown in Fig. 4B. In constant darkness (DD), neither *G2019S* nor *Rab10* manipulation affects the period. Exact genotypes in Table S4.

Fig. S8. **Rab 3 is found in a subset of dopaminergic neurons, differing from Rab10**. A. Rab3-GAL4 driven nuclear RFP (magenta) marks many neurons, including some that are dopaminergic (green). B. Coincidence is seen in the PAL neurons (B, white arrows), TH-VUM and DADN neurons (D, E), but not the MC neurons (C), where the green arrows indicate the MC neurons with no Rab3 staining, between other well-stained medulla neurons (magenta nuclei). Examination of the confocal images suggests that the PAM neurons

(B, green arrows) have no Rab3-driven RFP in their nuclei, though surrounding cells have well-marked nuclei. Summary of Rab3 expression in Fig. 5Cii. Exact genotypes in Table S5.

Fig. S9. **The phospho-Rab10 antibody specifically labels a few, non-dopaminergic neurons**. A. In wild-type flies, the phosopho-Rab10 antibody highlights a few neurons, in lateral and midline dorsal clusters, and in a ventral cluster. No dopamine neurons are intensely stained. Note that the ventral dopaminergic neurons (TH-VUM and DADN, yellow arrows) are not marked by phospho-Rab10. B. A second wild-type preparation, with only phospho-Rab10, indicates the same pattern of cells as A. C. In the *Rab10*$^{-}$ knock-out, no neurons are clearly stained (one weak cell body is indicated by a grey arrow). Maximum intensity projections of confocal stack through the fly brain, with B and C on same settings. Exact genotypes in Table S5.

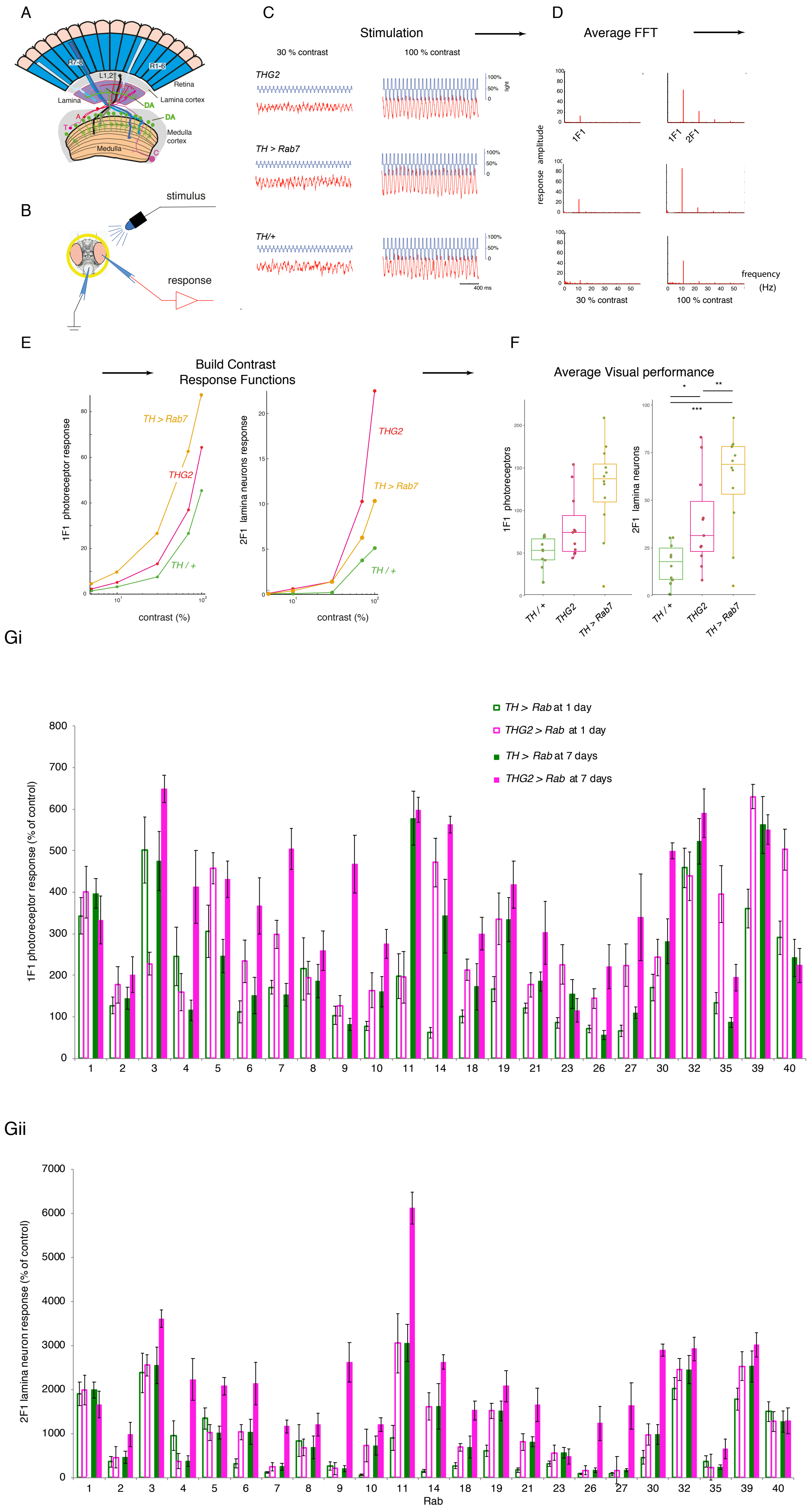
A
R7-8
R1-6
L1,2
Retina
Lamina
Lamina cortex
DA
DA
Medulla
Medulla cortex
B
stimulus
response
C
Stimulation
30 % contrast
100 % contrast
THG2
TH > Rab7
TH/+
100%
50%
0
light
400 ms
D
Average FFT
response amplitude
1F1
1F1 2F1
frequency (Hz)
30 % contrast
100 % contrast
E
Build Contrast Response Functions
1F1 photoreceptor response
2F1 lamina neurons response
TH > Rab7
THG2
TH / +
contrast (%)
F
Average Visual performance
1F1 photoreceptors
2F1 lamina neurons
TH / +
THG2
TH > Rab7
Gi
TH > Rab at 1 day
THG2 > Rab at 1 day
TH > Rab at 7 days
THG2 > Rab at 7 days
1F1 photoreceptor response (% of control)
1 2 3 4 5 6 7 8 9 10 11 14 18 19 21 23 26 27 30 32 35 39 40
Gii
2F1 lamina neuron response (% of control)
1 2 3 4 5 6 7 8 9 10 11 14 18 19 21 23 26 27 30 32 35 39 40
Rab

Fig S1

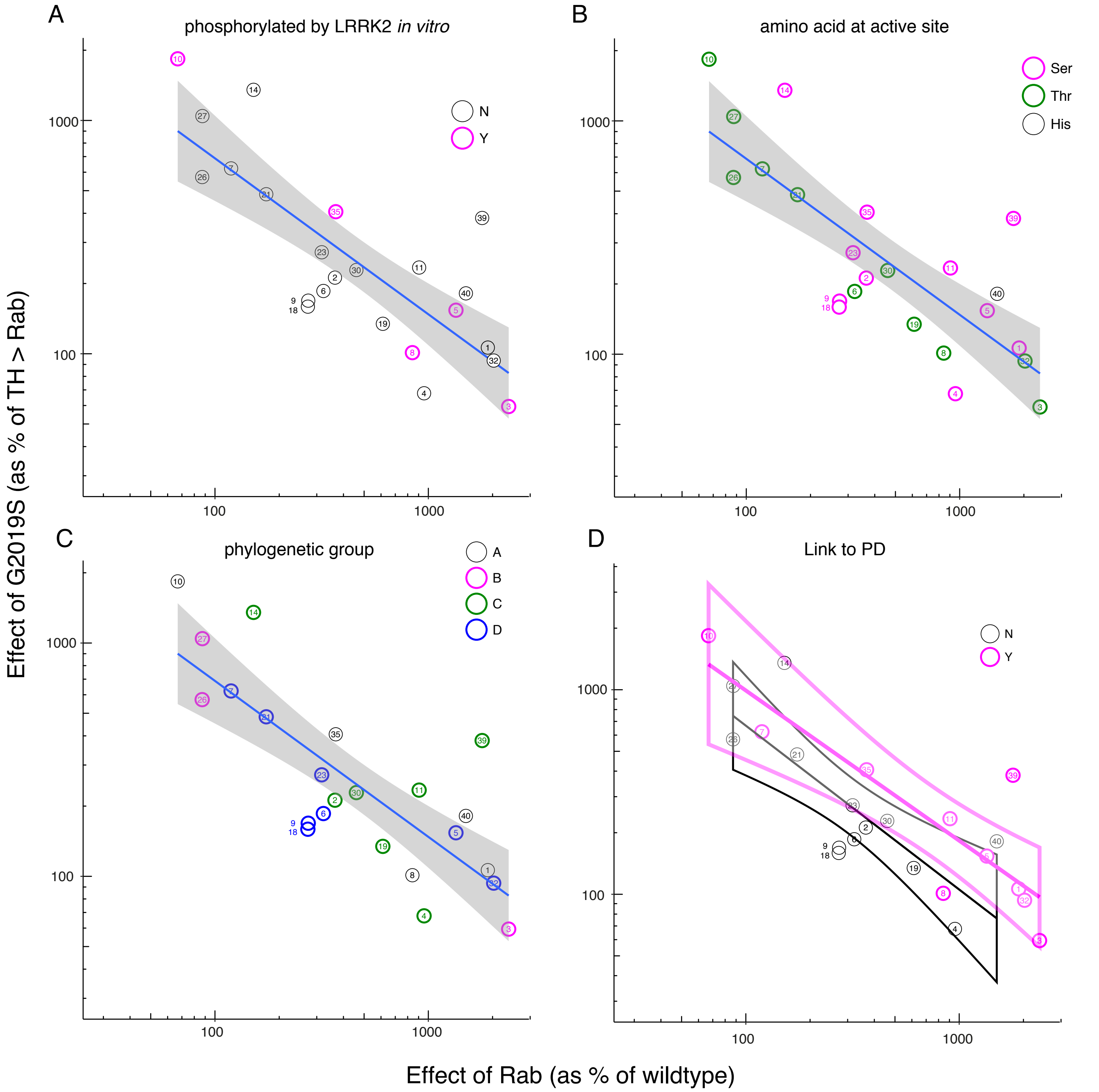
A
phosphorylated by LRRK2 in vitro
N
Y
B
amino acid at active site
Ser
Thr
His
C
phylogenetic group
A
B
C
D
D
Link to PD
N
Y
Effect of G2019S (as % of TH > Rab)
Effect of Rab (as % of wildtype)

Fig S2

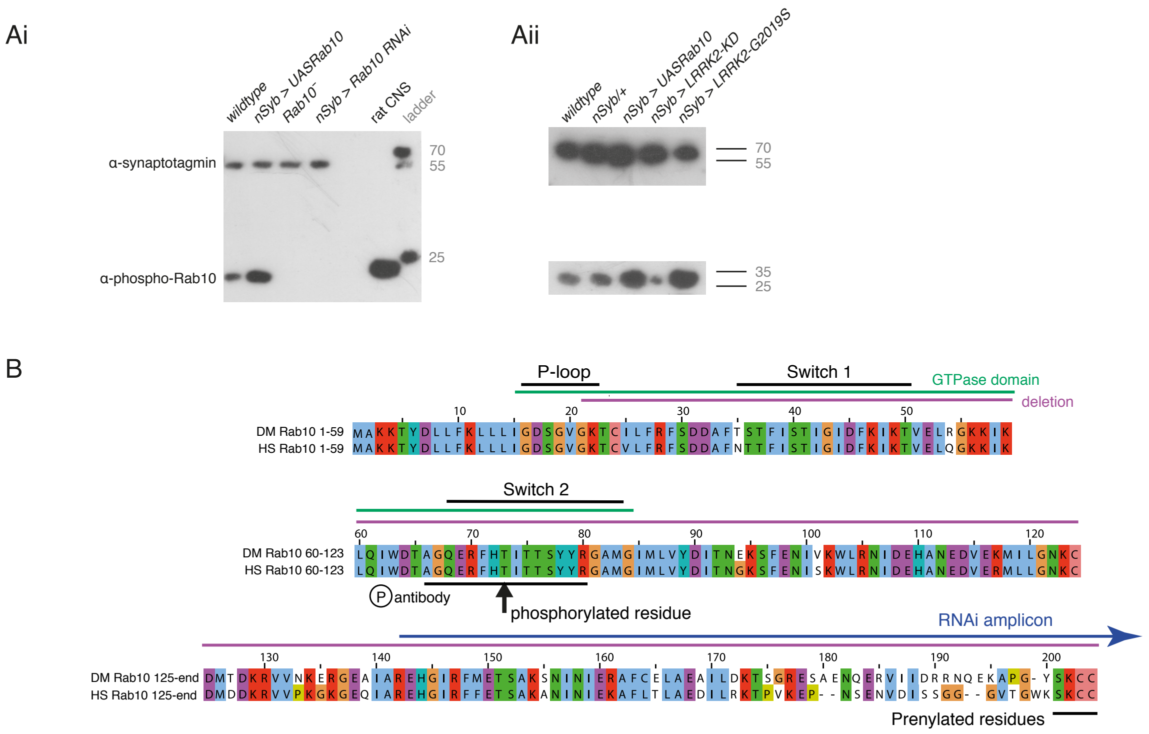
Ai
wildtype
nSyb > UASRab10
Rab10⁻
nSyb > Rab10 RNAi
rat CNS
ladder
α-synaptotagmin
α-phospho-Rab10
70
55
25
Aii
wildtype
nSyb/+
nSyb > UASRab10
nSyb > LRRK2-KD
nSyb > LRRK2-G2019S
70
55
35
25
B
P-loop
Switch 1
GTPase domain
deletion
DM Rab10 1-59
HS Rab10 1-59
Switch 2
DM Rab10 60-123
HS Rab10 60-123
antibody
phosphorylated residue
RNAi amplicon
DM Rab10 125-end
HS Rab10 125-end
Prenylated residues

Fig S3

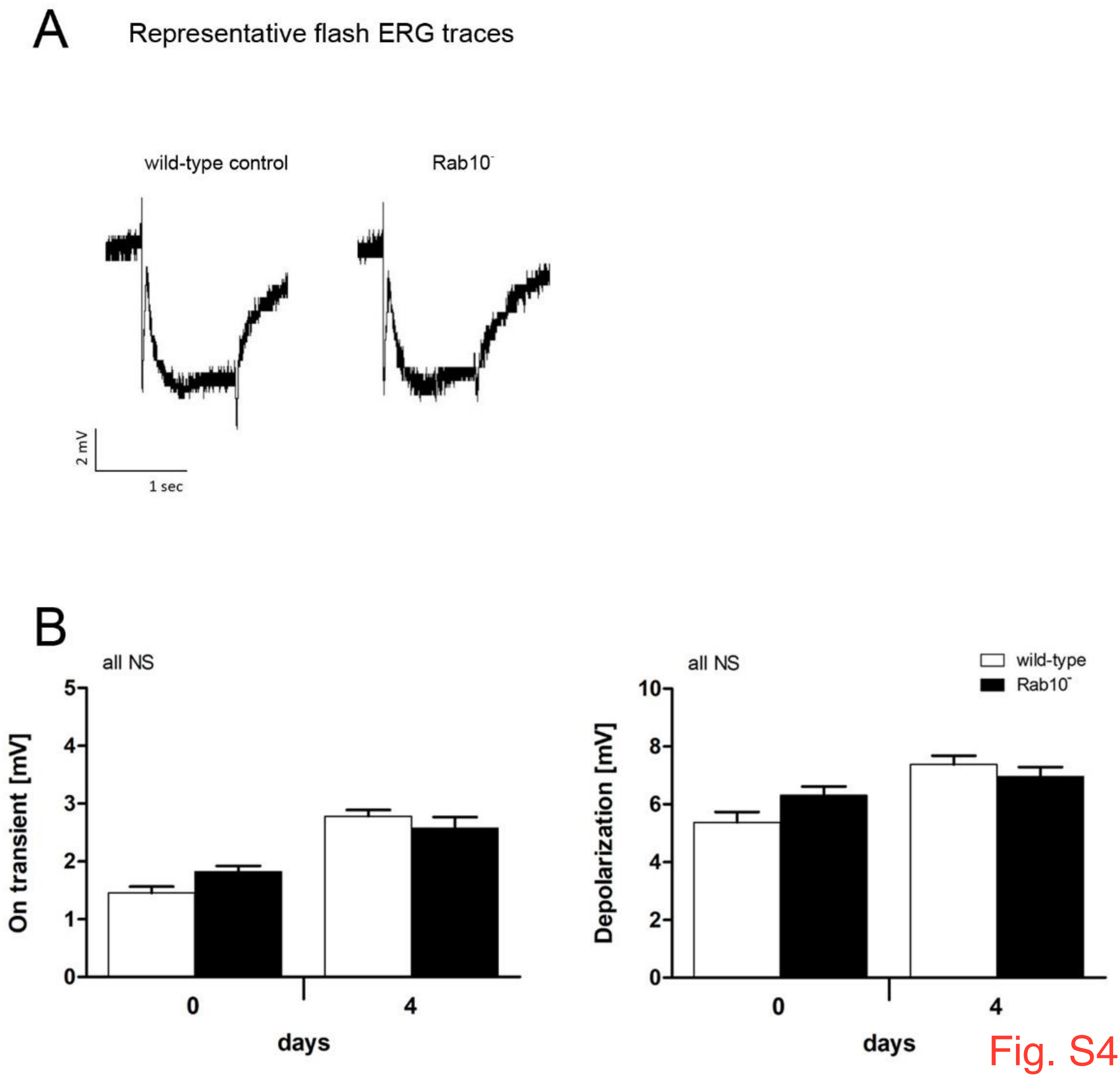
A
Representative flash ERG traces
wild-type control
Rab10⁻
2 mV
1 sec
B
all NS
On transient [mV]
5
4
3
2
1
0
0
4
days
all NS
wild-type
Rab10⁻
Depolarization [mV]
10
8
6
4
2
0
0
4
days

Fig. S4

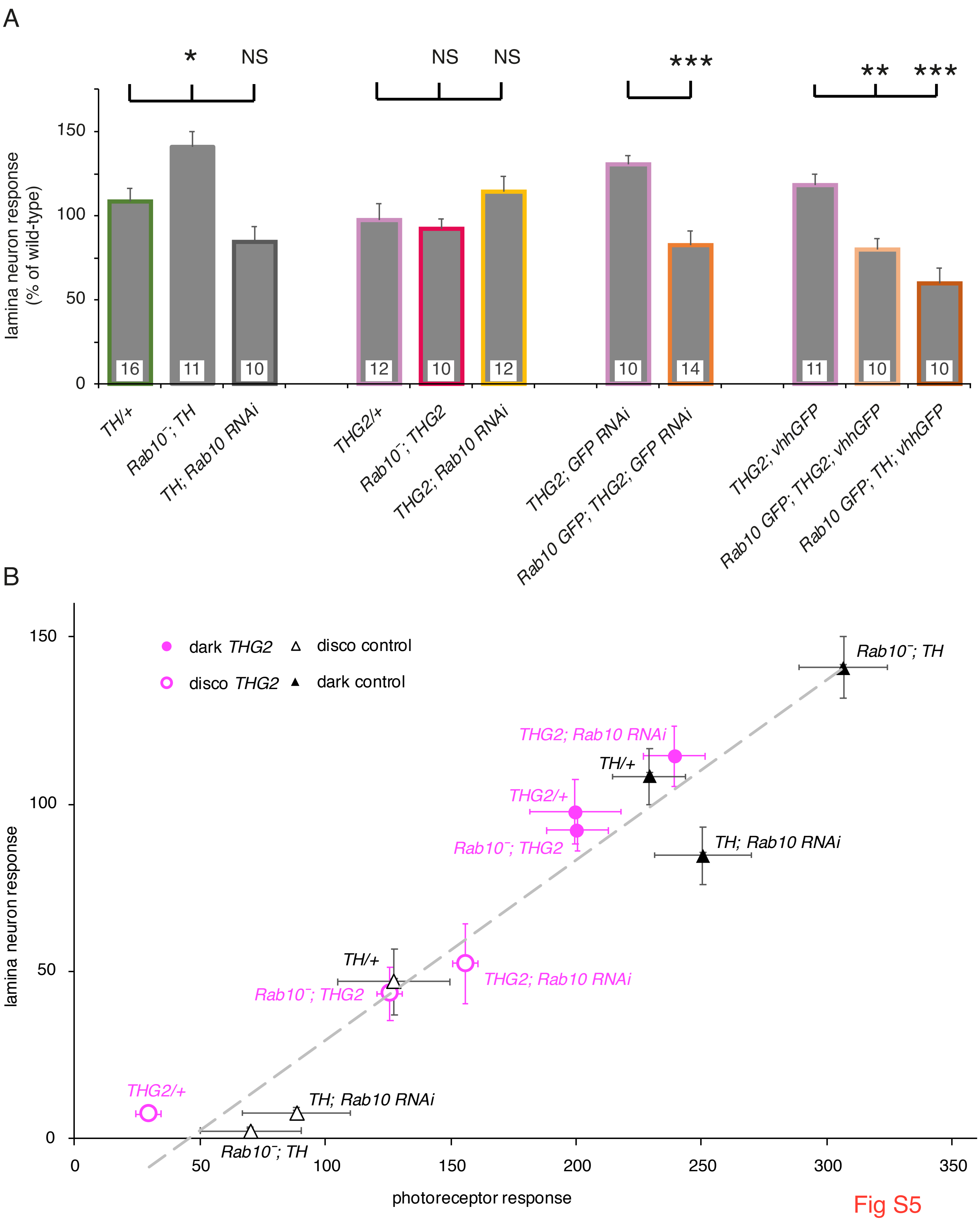
A
*
NS
NS
NS
***
**
***
lamina neuron response
(% of wild-type)
150
100
50
0
16
11
10
12
10
12
10
14
11
10
10
TH/+
Rab10⁻; TH
TH; Rab10 RNAi
THG2/+
Rab10⁻; THG2
THG2; Rab10 RNAi
THG2; GFP RNAi
Rab10 GFP; THG2; GFP RNAi
THG2; vhhGFP
Rab10 GFP; THG2; vhhGFP
Rab10 GFP; TH; vhhGFP
B
dark THG2
disco control
disco THG2
dark control
Rab10⁻; TH
THG2; Rab10 RNAi
TH/+
THG2/+
Rab10⁻; THG2
TH; Rab10 RNAi
TH/+
Rab10⁻; THG2
THG2; Rab10 RNAi
THG2/+
TH; Rab10 RNAi
Rab10⁻; TH
lamina neuron response
photoreceptor response
0
50
100
150
200
250
300
350

Fig S5

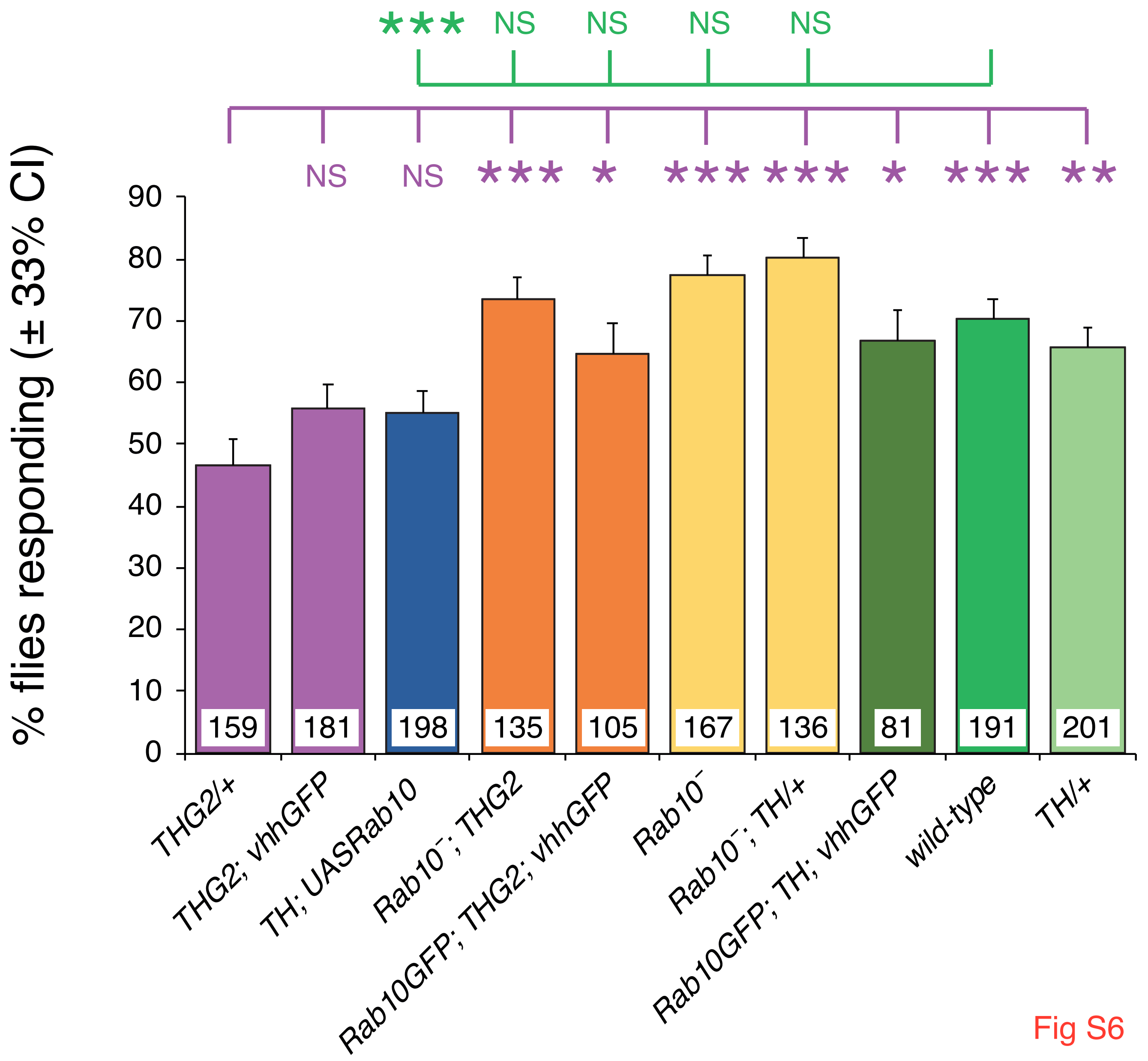

***
NS
NS
NS
NS
NS
NS
***
*
***
***
*
***
**
% flies responding (± 33% CI)
90
80
70
60
50
40
30
20
10
0
159
181
198
135
105
167
136
81
191
201
THG2/+
THG2; vhhGFP
TH; UASRab10
Rab10⁻; THG2
Rab10GFP; THG2; vhhGFP
Rab10⁻
Rab10⁻; TH/+
Rab10GFP; TH; vhhGFP
wild-type
TH/+

Fig S6

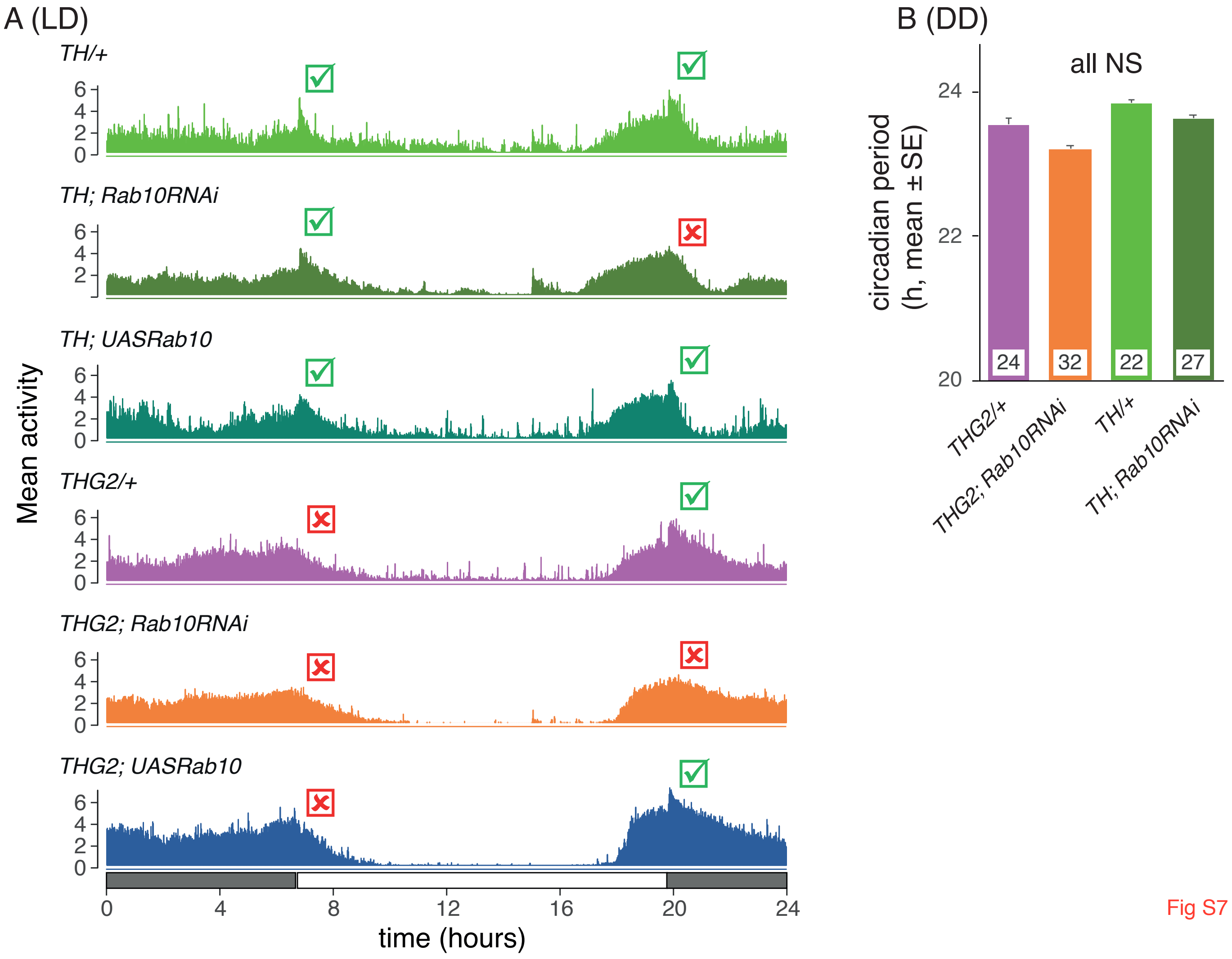

A (LD)
TH/+
TH; Rab10RNAi
TH; UASRab10
THG2/+
THG2; Rab10RNAi
THG2; UASRab10
Mean activity
0
2
4
6
0
4
8
12
16
20
24
time (hours)
B (DD)
all NS
circadian period
(h, mean ± SE)
20
22
24
24
32
22
27
THG2/+
THG2; Rab10RNAi
TH/+
TH; Rab10RNAi

Fig S7

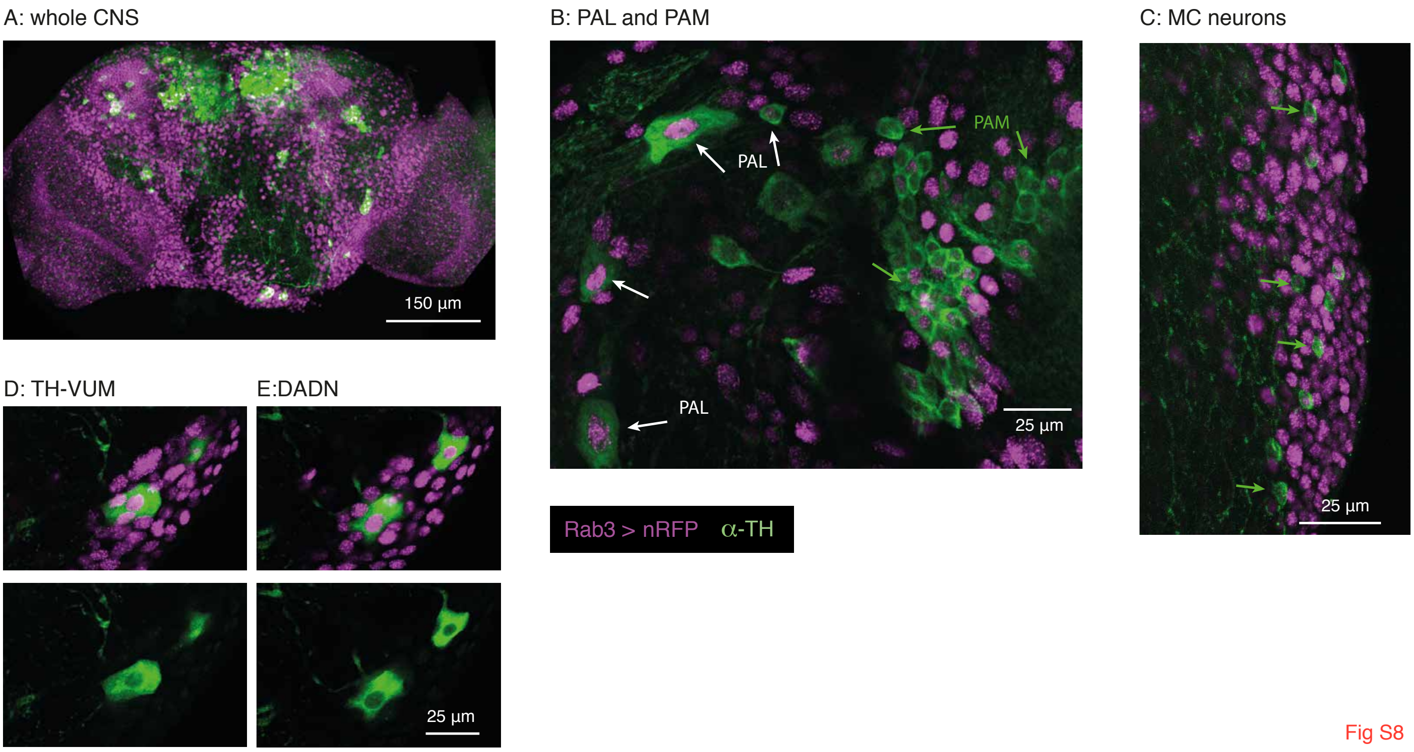
A: whole CNS
150 µm
B: PAL and PAM
PAL
PAM
PAL
25 µm
C: MC neurons
25 µm
D: TH-VUM
E:DADN
25 µm
Rab3 > nRFP
α-TH

Fig S8

**Supplementary Tables**

Table S1: Overexpression screen: exact genotypes and summary data: Mean and standard error of visual response of flies expressing RabGTPases with/without *LRRK2-G2019S*. Raw data for Fig. 1 and Fig. S2.

Table S2: Visual knockdown summary: genotypes and SSVEP responses for Fig. 2 and Fig. S5.

Table S3: Proboscis Extension Response Summary: genotypes and statistics for Fig. 3 and Fig. S6.

Table S4: Sleep Summary: genotypes and statistics for Fig.4 and Fig. S7.

Table S5: Anatomical Summary: genotypes and antibodies for Fig. 5, Fig. 8 and S9.

Table S6. List of fly Stocks.

Table S1 Mean and standard error of visual response of flies expressing RabGTPases with/without LRRK2-G2019S, as % of control (wildtype) flies
Raw data for Fig 1 and Fig. S2

1F1 photoreceptors

| | BDSC Stock number | 1F1 TH X Rab at 1 day | se | N | 1F1 TH X Rab at 7 day | se | N | 1F1 TH X Rab;G2109S at 1 day | se | N | 1F1 1 day increase due to G21019S | se | 1F1 TH X Rab;G2109S at 7 day | se | N | 1F1 7 day increase due to G21019S | se | |
|---|---|---|---|---|---|---|---|---|---|---|---|---|---|---|---|---|---|---|
| Rab1 | 24104 | 342.7 | 43.8 | 14 | 397.4 | 35.1 | 15 | 400.4 | 61.8 | 11 | 116.8 | 18.0 | 332.7 | 57.5 | 12 | 83.7 | 14.4 | |
| Rab2 | 23246 | 127.6 | 19.9 | 10 | 144.8 | 26.5 | 12 | 177.5 | 43.9 | 9 | 139.1 | 34.4 | 201.6 | 42.2 | 9 | 139.3 | 23.8 | |
| Rab3 | 9762 | 501.4 | 79.8 | 10 | 474.8 | 71.7 | 10 | 227.4 | 27.3 | 10 | 45.4 | 5.5 | 648.6 | 33.2 | 10 | 136.6 | 14.6 | |
| Rab4 | 9767 | 245.0 | 70.1 | 10 | 115.9 | 25.3 | 13 | 159.7 | 44.8 | 10 | 65.2 | 18.3 | 413.5 | 87.9 | 9 | 356.9 | 55.0 | |
| Rab5 | 50788 | 305.2 | 63.0 | 11 | 247.0 | 40.0 | 11 | 458.4 | 37.6 | 12 | 150.2 | 12.3 | 431.3 | 43.1 | 14 | 174.6 | 9.4 | |
| Rab6 | 23251 | 111.7 | 26.0 | 11 | 152.0 | 44.1 | 10 | 234.1 | 51.0 | 10 | 209.6 | 45.7 | 367.9 | 67.6 | 9 | 242.1 | 28.9 | |
| Rab7 | 23270 | 171.3 | 16.9 | 11 | 153.0 | 26.9 | 10 | 298.5 | 34.1 | 12 | 174.3 | 19.9 | 504.2 | 49.9 | 10 | 329.5 | 16.7 | |
| Rab8 | 9782 | 215.8 | 74.7 | 10 | 185.8 | 40.5 | 12 | 194.3 | 38.7 | 10 | 90.0 | 17.9 | 259.6 | 47.1 | 11 | 139.7 | 24.2 | |
| Rab9 | 9784 | 103.5 | 22.4 | 12 | 81.7 | 15.0 | 10 | 127.2 | 25.0 | 9 | 122.8 | 24.2 | 467.5 | 70.0 | 9 | 571.9 | 55.1 | |
| Rab10 | 24097 | 77.9 | 11.4 | 10 | 160.5 | 37.4 | 10 | 163.8 | 42.5 | 11 | 210.2 | 54.5 | 276.2 | 34.5 | 10 | 172.1 | 21.1 | |
| Rab11 | 50782 | 197.7 | 53.5 | 10 | 578.2 | 65.2 | 10 | 195.9 | 62.3 | 10 | 99.1 | 31.5 | 598.6 | 29.5 | 10 | 103.5 | 15.0 | |
| Rab14 | 9794 | 61.8 | 12.7 | 10 | 343.0 | 88.5 | 10 | 471.6 | 59.3 | 10 | 763.0 | 96.0 | 562.8 | 20.4 | 10 | 164.1 | 4.3 | |
| Rab18 | 9796 | 100.9 | 15.5 | 10 | 173.1 | 55.7 | 10 | 213.5 | 26.3 | 15 | 211.6 | 26.1 | 300.4 | 39.4 | 22 | 173.5 | 18.4 | |
| Rab19 | 24150 | 167.0 | 30.3 | 12 | 334.7 | 53.1 | 11 | 335.9 | 62.4 | 9 | 201.2 | 37.4 | 418.1 | 56.6 | 10 | 124.9 | 16.8 | |
| Rab21 | 23242 | 120.8 | 12.1 | 9 | 185.6 | 22.6 | 11 | 177.3 | 29.9 | 9 | 146.8 | 24.7 | 302.4 | 75.4 | 10 | 162.9 | 42.5 | |
| Rab23 | 9802 | 85.7 | 13.4 | 12 | 154.5 | 34.9 | 13 | 226.2 | 48.0 | 9 | 263.9 | 56.0 | 115.5 | 28.9 | 11 | 74.7 | 12.8 | |
| Rab26 | 23245 | 71.5 | 9.4 | 12 | 55.7 | 10.8 | 13 | 144.1 | 24.1 | 9 | 201.7 | 33.7 | 221.1 | 52.9 | 10 | 396.9 | 36.7 | |
| Rab27 | 9810 | 66.0 | 13.7 | 13 | 110.2 | 13.6 | 10 | 224.5 | 51.3 | 10 | 340.1 | 77.7 | 339.0 | 104.3 | 10 | 307.7 | 46.4 | |
| Rab30 | 9812 | 170.0 | 31.7 | 10 | 281.8 | 54.0 | 10 | 243.0 | 44.2 | 9 | 142.9 | 26.0 | 499.6 | 19.2 | 10 | 177.3 | 7.9 | |
| Rab32 | 9815 | 458.9 | 47.1 | 10 | 522.6 | 55.3 | 14 | 438.7 | 58.3 | 10 | 95.6 | 12.7 | 590.3 | 58.2 | 10 | 307.4 | 13.3 | |
| Rab35 | 9821 | 133.2 | 26.2 | 12 | 86.8 | 12.0 | 15 | 395.9 | 67.9 | 10 | 297.1 | 51.0 | 194.5 | 32.4 | 11 | 224.2 | 8.2 | |
| Rab39 | 9825 | 360.8 | 45.9 | 10 | 562.4 | 69.3 | 10 | 630.7 | 30.1 | 10 | 174.8 | 8.3 | 551.0 | 35.7 | 10 | 98.0 | 5.7 | |
| Rab40 | 9830 | 290.5 | 41.0 | 12 | 242.4 | 44.6 | 11 | 502.7 | 48.8 | 11 | 173.1 | 16.8 | 223.7 | 41.6 | 12 | 92.3 | 8.3 | |
| total flies | | | | 251 | | | 261 | | | 235 | | | | | 249 | Grand total | | 996 |

| Fig. S2A Phosphorylated in vitro (Steger et al, 2017) | Fig. S2B. Amino acid at active site (Zhang et al, 2007) | Fig. S2C. Phylogenetic group per Zhang et al (2007) | Fig. S2D. Previously linked to Parkinson's (Shi et al, 201 | |
|---|---|---|---|---|
| N | S | A | Y | Rab1 |
| N | S | C | N | Rab2 |
| Y | T | B | Y | Rab3 |
| N | S | C | N | Rab4 |
| Y | S | D | Y | Rab5 |
| N | T | D | N | Rab6 |
| N | T | D | Y | Rab7 |
| Y | T | A | Y | Rab8 |
| N | S | D | N | Rab9 |
| Y | T | A | Y | Rab10 |
| N | S | C | Y | Rab11 |
| N | S | C | N | Rab14 |
| N | S | D | N | Rab18 |
| N | T | C | N | Rab19 |
| N | T | D | N | Rab21 |
| N | S | D | N | Rab23 |
| N | T | B | N | Rab26 |
| N | T | B | N | Rab27 |
| N | T | C | N | Rab30 |
| | T | D | Y | Rab32 |
| Y | S | A | Y | Rab35 |
| N | S | C | Y | Rab39 |
| N | H | A | N | Rab40 |

2F1 lamina neurons

| | | | 1F1 TH X Rab at 1 day | se | N | | 1F1 TH X Rab at 7 day | se | N | | 1F1 TH X Rab;G2109S at 1 day | se | N | | 1F1 1 day increase due to G21019[illegible] | se | | 1F1 TH X Rab;G2109S at 7 day | se | N | | 1F1 7 day increase due to G21019[illegible] | se |
|---|---|---|---|---|---|---|---|---|---|---|---|---|---|---|---|---|---|---|---|---|---|---|---|
| Rab1 | | | 1900.1 | 267.8 | 14 | | 1994.7 | 182.4 | 15 | | 2019.1 | 337.5 | 11 | | 106.3 | 17.8 | | 1657.5 | 298.6 | 12 | | 83.1 | 14.8 |
| Rab2 | | | 365.1 | 119.1 | 10 | | 461.3 | 133.9 | 12 | | 773.9 | 241.4 | 9 | | 212.0 | 66.1 | | 977.8 | 273.9 | 9 | | 212.0 | 35.4 |
| Rab3 | | | 2379.6 | 442.4 | 10 | | 2549.9 | 418.8 | 11 | | 1413.5 | 244.6 | 11 | | 59.4 | 10.3 | | 3607.0 | 198.2 | 10 | | 141.5 | 14.0 |
| Rab4 | | | 955.0 | 342.9 | 10 | | 373.6 | 117.9 | 13 | | 646.9 | 176.8 | 10 | | 67.7 | 18.5 | | 2229.4 | 475.5 | 9 | | 596.8 | 73.5 |
| Rab5 | | | 1347.0 | 241.8 | 11 | | 1020.5 | 145.9 | 11 | | 2070.2 | 179.8 | 12 | | 153.7 | 13.3 | | 2079.8 | 196.1 | 14 | | 203.8 | 9.5 |
| Rab6 | | | 322.2 | 100.5 | 11 | | 1037.6 | 284.8 | 13 | | 598.6 | 171.6 | 7 | | 185.8 | 53.3 | | 2137.5 | 484.4 | 9 | | 206.0 | 80.9 |
| Rab7 | | | 119.0 | 19.0 | 11 | | 248.2 | 71.4 | 10 | | 741.5 | 87.5 | 12 | | 623.0 | 73.5 | | 1164.7 | 150.2 | 10 | | 469.2 | 20.3 |
| Rab8 | | | 842.0 | 360.5 | 10 | | 685.9 | 256.6 | 12 | | 851.4 | 195.8 | 10 | | 101.1 | 23.3 | | 1201.2 | 259.8 | 11 | | 175.1 | 30.5 |
| Rab9 | | | 273.6 | 95.0 | 12 | | 207.7 | 67.8 | 10 | | 462.2 | 138.4 | 9 | | 168.9 | 50.6 | | 2620.3 | 448.7 | 9 | | 1261.5 | 97.1 |
| Rab10 | | | 66.9 | 10.2 | 10 | | 730.4 | 220.1 | 10 | | 1228.7 | 368.4 | 14 | | 1837.5 | 550.9 | | 1206.6 | 152.1 | 13 | | 165.2 | 12.4 |
| Rab11 | | | 905.4 | 288.6 | 10 | | 3052.9 | 420.5 | 10 | | 2120.5 | 674.3 | 10 | | 234.2 | 74.5 | | 6117.6 | 365.7 | 10 | | 200.4 | 17.2 |
| Rab14 | | | 151.6 | 40.5 | 10 | | 1618.1 | 522.3 | 10 | | 2051.2 | 308.4 | 10 | | 1352.9 | 203.4 | | 2625.0 | 166.9 | 10 | | 162.2 | 8.1 |
| Rab18 | | | 272.5 | 71.9 | 10 | | 691.8 | 250.7 | 10 | | 433.9 | 83.4 | 15 | | 159.2 | 30.6 | | 1530.2 | 215.3 | 22 | | 221.2 | 49.6 |
| Rab19 | | | 612.0 | 133.7 | 12 | | 1516.4 | 218.6 | 11 | | 822.7 | 178.9 | 6 | | 134.4 | 29.2 | | 2076.0 | 352.4 | 10 | | 136.9 | 42.8 |
| Rab21 | | | 173.9 | 56.5 | 7 | | 811.5 | 111.9 | 11 | | 839.8 | 194.0 | 9 | | 482.8 | 111.6 | | 1649.9 | 382.1 | 10 | | 203.3 | 45.5 |
| Rab23 | | | 316.5 | 66.8 | 12 | | 563.5 | 116.1 | 13 | | 861.8 | 168.2 | 9 | | 272.3 | 53.2 | | 478.9 | 169.1 | 11 | | 85.0 | 19.6 |
| Rab26 | | | 87.1 | 13.6 | 12 | | 169.1 | 51.6 | 13 | | 498.1 | 110.0 | 9 | | 572.0 | 126.3 | | 1235.3 | 388.1 | 10 | | 730.4 | 77.9 |
| Rab27 | | | 87.3 | 30.1 | 13 | | 169.1 | 29.6 | 10 | | 911.5 | 308.2 | 10 | | 1044.1 | 353.0 | | 1628.8 | 524.6 | 10 | | 963.2 | 57.6 |
| Rab30 | | | 460.0 | 157.6 | 8 | | 978.3 | 225.2 | 10 | | 1051.2 | 240.7 | 9 | | 228.5 | 52.3 | | 2893.5 | 138.9 | 10 | | 295.8 | 13.2 |
| Rab32 | | | 2024.4 | 251.7 | 10 | | 2452.2 | 318.9 | 14 | | 1893.6 | 250.1 | 10 | | 93.5 | 12.4 | | 2925.0 | 268.7 | 10 | | 119.3 | 14.2 |
| Rab35 | | | 368.2 | 128.7 | 12 | | 238.6 | 48.7 | 15 | | 1495.8 | 297.6 | 10 | | 406.3 | 80.8 | | 654.7 | 218.1 | 7 | | 274.4 | 14.6 |
| Rab39 | | | 1787.4 | 248.5 | 10 | | 2530.0 | 355.3 | 10 | | 6825.5 | 326.0 | 10 | | 381.9 | 18.2 | | 3017.7 | 271.9 | 10 | | 119.3 | 4.0 |
| Rab40 | | | 1499.5 | 217.3 | 12 | | 1275.6 | 240.6 | 11 | | 2721.2 | 228.3 | 11 | | 181.5 | 15.2 | | 1290.7 | 295.1 | 12 | | 101.2 | 10.8 |

Table S2. Summary table of genotypes and SSVEP responses for Fig. 2 and Fig. S5

| | | Lamina neuron response | | | | | | | | Photoreceptor response | | | | | | |
|---|---|---|---|---|---|---|---|---|---|---|---|---|---|---|---|---|
| | | dark | | | | disco | | | | dark | | | | disco | | |
| Label | Formal genotype | mean | N | SE | | mean | N | SE | | mean | N | SE | | mean | N | SE |
| *TH/+* | *+; +; TH/w1118* | 108.3 | 16 | 8.3 | | 46.9 | 12 | 10.0 | | 229.2 | 17 | 14.6 | | 127.4 | 12 | 22.4 |
| *Rab10*$^{-}$*; TH* | *Rab10*$^{-}$*; +; TH/+* | 140.8 | 11 | 9.2 | | 2.4 | 8 | 0.5 | | 306.5 | 11 | 17.7 | | 70.0 | 9 | 20.2 |
| *TH; Rab10 RNAi* | *+; dicer2/+; TH/Rab10 RNAi 26289* | 84.7 | 10 | 8.6 | | 7.6 | 10 | 1.7 | | 250.5 | 10 | 19.2 | | 88.5 | 13 | 21.6 |
| *THG2/+* | *+; +; TH::LRRK2-G2019S/w1118* | 97.6 | 12 | 9.6 | | 7.4 | 11 | 1.6 | | 199.6 | 13 | 18.2 | | 29.5 | 11 | 7.8 |
| *Rab10*$^{-}$*; THG2* | *Rab10*$^{-}$*; +; TH::LRRK2-G2019S/+* | 92.1 | 10 | 6.2 | | 43.3 | 12 | 8.1 | | 200.5 | 10 | 12.2 | | 125.8 | 12 | 24.0 |
| *THG2; Rab10 RNAi* | *+; dicer2/+; TH::LRRK2-G2019S/Rab10 RNAi 26289* | 114.3 | 12 | 8.8 | | 52.3 | 12 | 12.0 | | 239.1 | 12 | 12.3 | | 155.8 | 12 | 28.6 |
| *THG2; GFP RNAi* | *+; GFP RNAi 9331/+; TH::LRRK2-G2019S/+* | 130.7 | 10 | 4.8 | | 11.8 | 9 | 3.0 | | 277.9 | 10 | 8.7 | | 48.9 | 9 | 12.7 |
| *Rab10 GFP; THG2; GFP RNAi* | *Rab10 GFP; GFP RNAi 9331/+; TH::LRRK2-G2019S/+* | 82.4 | 14 | 8.3 | | 61.4 | 12 | 11.8 | | 192.5 | 14 | 13.5 | | 199.8 | 12 | 25.9 |
| *THG2; vhhGFP* | *+; P-NSlmb-vhhGFP/+; TH::LRRK2-G2019S/+* | 118.0 | 11 | 6.5 | | 1.0 | 11 | 0.2 | | 272.0 | 11 | 6.9 | | 8.3 | 11 | 1.9 |
| *Rab10 GFP; THG2; vhhGFP* | *Rab10 GFP; P-NSlmb-vhhGFP/+; TH::LRRK2-G2019S/+* | 80.1 | 10 | 6.4 | | 17.8 | 11 | 4.3 | | 225.5 | 10 | 17.6 | | 78.4 | 11 | 19.7 |
| *Rab10 GFP; TH; vhhGFP* | *Rab10 GFP; P-NSlmb-vhhGFP/+; TH/+* | 60.1 | 10 | 9.0 | | 13.8 | 11 | 3.6 | | 211.3 | 10 | 19.8 | | 77.9 | 11 | 18.8 |

Table S3. Summary data for Figs. 3 and S6

Fig. 3

| Formal Genotype | Label |
|---|---|
| *Gr5a-Lex::LexOp-ReachR/+;+/+* | *Gr5a>ReachR* |
| *Gr5a-Lex::LexOp-ReachR/+;TH::LRRK2-G2019S/+* | *THG2; Gr5a>ReachR* |
| *Gr5a-Lex::LexOp-ReachR/dicer2;TH::LRRK2-G2019S/Rab10 RNAi 26289* | *THG2; Rab10RNAi; Gr5a>ReachR* |

Fig. S6

Akinesia: $\chi^2$ test

Overall $\chi^2$ = 70.3, df = 9, p-value = 1.3e-11

Post hoc P values from 'fifer' package in R

| Formal Genotype | Label | *THG2; vhhGFP* | *TH; UASRab10* | *Rab10$^{-}$; THG2* | *Rab10GFP; THG2>vhhGFP* | *Rab10$^{-}$* | *Rab10$^{-}$; TH/+* | *Rab10GFP; TH>vhhGFP* | *wild-type* | TH/+ |
|---|---|---|---|---|---|---|---|---|---|---|
| *+; +; TH::LRRK2-G2019S/w1118* | ***THG2/+*** | **0.1744** | **0.1744** | **0** | **0.0128** | **0** | **0** | **0.0128** | **0.0001** | **0.0019** |
| *+; P-NSlmb-vhhGFP/+; TH::LRRK2-G2019S/w1118* | *THG2; vhhGFP* | | 0.9178 | 0.006 | 0.2316 | 0.0001 | 0 | 0.1744 | **0.0144** | 0.1145 |
| *TH > UAS Rab10 24097* | *TH; UASRab10* | | | 0.0037 | 0.1744 | 0.0001 | 0 | 0.1551 | **0.0089** | 0.072 |
| *Rab10$^{-}$; +; TH::LRRK2-G2019S/+* | *Rab10$^{-}$; THG2* | | | | 0.2252 | 0.5937 | 0.2621 | 0.4532 | **0.679** | 0.2189 |
| *Rab10 GFP; P-NSlmb-vhhGFP/+; TH::LRRK2-G2019S/+* | *Rab10GFP; THG2>vhhGFP* | | | | | 0.0633 | 0.022 | 0.9178 | **0.4532** | 0.9178 |
| *Rab10$^{-}$; +; TH/+* | *Rab10$^{-}$* | | | | | | 0.6476 | 0.1629 | **0.2189** | 0.0394 |
| *Rab10$^{-}$; +; + [Rab10$^{-}$ ♀□ x ♂□ w1118]* | *Rab10$^{-}$; TH/+* | | | | | | | 0.0739 | **0.1099** | 0.0137 |
| *Rab10 GFP; P-NSlmb-vhhGFP/+; TH/+* | *Rab10GFP; TH>vhhGFP* | | | | | | | | **0.6476** | 0.9178 |
| *+/+ [CS x w1118]* | ***wild-type*** | | | | | | | | | **0.4706** |
| *+ +; TH/w1118* | *TH/+* | | | | | | | | | |

### Table S4. Summary data and genotypes for sleep and circadian data

Fig. 4 and S7

| | | Sleep | | | | | | | Circadian | | |
|---|---|---|---|---|---|---|---|---|---|---|---|
| | | Sleep in 12:12 LD | | | | | | | DD | | |
| | | Day | | | | Night | | | circadian period (hours) | | |
| Formal genotype | label | N | mean | se | | N | mean | se | N | mean | se |
| *+; +; TH/ev (empty RNAi vector 36303)* | *TH/+* | 15 | 0.563 | 0.022 | 0 | 15 | 0.355 | 0.046 | 22 | 23.8 | 0.05 |
| *+; +; TH::LRRK2-G2019S/ev (empty RNAi vector 36303)* | *THG2/+* | 15 | 0.647 | 0.028 | 0 | 15 | 0.129 | 0.021 | 24 | 23.6 | 0.09 |
| *+; dicer2/+; TH::LRRK2-G2019S/Rab10 RNAi 26289* | *THG2; Rab10 RNAi* | 16 | 0.757 | 0.022 | 0 | 16 | 0.118 | 0.025 | 32 | 23.2 | 0.04 |
| *+; RAB10/+; TH/+* | *TH; UASRab10* | 16 | 0.607 | 0.021 | 0 | 16 | 0.433 | 0.030 | Not determined | | |
| *+; dicer2/+; TH/Rab10 RNAi 26289* | *TH; Rab10 RNAi* | 14 | 0.594 | 0.021 | 0 | 14 | 0.386 | 0.039 | 27 | 23.6 | 0.05 |
| *+; RAB10/+; TH::LRRK2-G2019S/+* | *THG2; UASRab10* | 16 | 0.756 | 0.017 | 0 | 16 | 0.121 | 0.032 | Not determined | | |

Table S5 Genotypes for Fig. 5, S8 and Fig. S9

Fig. 5

| | Genotype | Antibody |
|---|---|---|
| A i-A iii | *+; RedStinger4 nRFP/+; Rab10 Gal4/+* | Mouse anti TH Immunostar (22941, 1:1000) |
| A iv | *+; TH-LexA/UAS-CD4-spGFP1-10; Rab10-Gal4/lexAop-CD4-spGFP11* | Mouse anti TH Immunostar (22941, 1:1000) |

Fig. S8

| | | |
|---|---|---|
| A,B,C | *+; RedStinger4 nRFP/+; Rab3 Gal4/+* | Mouse anti TH Immunostar (22941, 1:1000) |

Fig. S9

| | | |
|---|---|---|
| A | *w1118* | Mouse anti TH Immunostar (22941, 1:1000) and phospho-Rab10 (Abcam: ab230261, 1:250) |
| B | *w1118* | phospho-Rab10 (Abcam: ab230261, 1:250) |
| C | *Rab10*$^{-}$ | phospho-Rab10 (Abcam: ab230261, 1:250) |

Supplementary Table S6. List of Fly stocks

| Genotype | kind gift of |
|---|---|
| *CS (Canton-S)* | Sean Sweeney |
| *Gr5a-LexA* | Kristin Scott |
| *LRRK2-G2019S* | Wanli Smith |
| *LRRK2-G2019S-K1906M (LRRK2-KD)* | Cheng-Ting Chien |
| *nSyb-GAL4* | Julie Simpson |
| *Rab10⁻* | this paper |
| *TH-GAl4* | Serge Birman |
| *TH-LexA* | Yoshi Aso |
| *UAS-CD4-spGFP1-10; LexAop-CD4-spGFP11* | Mattias Landgraf |
| *vhhGFP* | Stefan Heidmann |
| *w1118 (w⁻)* | Sean Sweeney |

| Genotype | Bloomington Stock |
|---|---|
| *Dicer2* | 24650 |
| *LexOp-ReachR* | 53747 |
| *GFP RNAi* | 9331 |
| *nRFP* | 8546 |
| *Rab10 GAL4* | 51588 |
| *Rab10 GFP (TI{TI}Rab10 EYFP)* | 62548 |
| *Rab10 RNAi* | 26289 |

| UAS-Rab stocks are from Bloomington and are listed in Table S1 |
|---|
